# Tuft dendrite spikes are accompanied by selective input from distinct functional networks

**DOI:** 10.64898/2026.06.10.731464

**Authors:** Jacob Gable, Zachary L Newman, Sarah Young, Jackson Scheib, Savannah Bliese, Nicole Miller, Aaron Kerlin

## Abstract

The tuft dendrites of layer 5 neurons can support regenerative events – dendritic spikes – that have been proposed to coordinate context-specific engagement and plasticity within cortical networks. However, it remains unclear whether tuft spikes are accompanied by input activity with dynamics that could support these network-level functions. To address this, glutamatergic synapses and postsynaptic calcium signals were simultaneously imaged in the tuft dendrites of layer 5 extratelencephalic neurons within the premotor cortex of mice performing a cued directional licking task. Trial-to-trial, the generation of tuft spikes was associated with a multiphasic elevation in synaptic activity spanning hundreds of milliseconds. This activity was highly specific to the dendrite in which a spike was detected, suggesting the concurrent activation of select subnetworks. Synapses that were strongly coupled to the overall population were the most synchronized with tuft spikes and preferentially encoded the transition between the preparation and action epochs of the task. Even among these strongly coupled synapses, increases in activity were largely specific to synapses located on the spiking dendrite. Surprisingly, among synapses with the poorest population coupling, a second population of coactive synapses was discovered that was also associated with tuft spikes and functionally selective for task-outcome. These results suggest that tuft spikes may be particularly driven by inputs from neurons that are both embedded in sparse subnetworks and synchronized through coupling to larger-scale functional networks.

**Significance Statement:** Flexible behavior and learning may depend on interactions between activity in different brain networks and dendritic spikes generated within the neurons that make up the output layer of the neocortex. The results of this study indicate that at the moment of spike generation, spikes in different dendrites are associated with the activation of very specific networks. Yet, across time, the inputs most associated with dendritic spikes are broadly coactive and share selectivity for similar features of behavior. This suggests that dendritic spikes may be particularly driven by the activation of “hub” neurons that coordinate communication between large-scale and small-scale functional networks.

## Introduction

Neurons do not simply compute the sum of all their inputs. This is especially apparent for layer 5 (L5) extratelencephalic (ET) cortical neurons, the primary output channel of the cortex. Their apical tuft dendrites elaborate in layer 1, far from the cell soma and basal dendrites. The apical tuft dendrites contain active conductances that can support powerful regenerative events – dendritic spikes – triggered by nonlinear integration of synaptic inputs with or without back-propagating action potentials (bAPs; 1–4). These tuft spikes can trigger large calcium influx, prolonged somatic depolarization, and burst firing (1). Evidence suggests that tuft spikes condition the output of L5 neurons on long-range contextual signals (5–8), and rapidly induce long-lasting functional changes (9–11).

To understand the patterns of synaptic inputs that trigger these sparse events, tuft dendrites have been extensively studied in *ex vivo* brain tissue, where input patterns can be precisely controlled. Short-timescale (≈ 10 ms) co-activation of spatially clustered inputs can be especially effective at engaging dendritic nonlinearities (3, 12–15), consistent with the idea that small segments of dendrite could operate as independent computational subunits (16), dramatically increasing the input pattern discrimination (17) and information storage (18) capabilities of individual neurons. In support of this model, *in vivo* functional imaging of the tuft indicates that inputs with similar function and trial-to-trial coactivity tend to cluster together (19). However, *in vivo* imaging also indicates that fully isolated spiking activity within small tuft segments is rare (19–23) and that only larger-scale activity within the apical tree is reliably associated with somatic output (23). This large-scale activity could still be dependent on clustered inputs (23), but the paucity of fully independent tuft subunits *in vivo* suggests that dispersed input coactive over longer timescales may play a prominent role in tuft computation.

Recent models have focused on a potential role for dendritic spikes in more network-level operations (24–29). These models imply that dendritic spikes occur within specific input episodes that reflect coordinated activation of behavior-related subnetworks at both long and short timescales. For example, evidence indicates that dendritic plateau potentials trigger the formation of new place fields in hippocampal neurons (9) based on long timescale input activity, suggesting a process operating at the population-level rather than pairwise correlations (29). Tuft spikes have also been proposed to trigger recurrent activity that coordinates learning across subnetworks that span the cortical hierarchy (28). In addition to these roles in coordinating plasticity, dendritic spikes have been proposed to enable context-dependent computation by gating cortical hypernetworks (30, 31) or by facilitating synchronous firing of different functional subnetworks engaged in the same task (26). All of these models predict that spikes in distal dendrites should be accompanied by subnetwork-specific input activity to that dendrite at both long and short timescales. Yet, little is known about the dynamics of glutamatergic input to the tuft that accompanies dendritic spikes *in vivo* or its relationship to the functional subnetworks engaged during behavior.

Here, we simultaneously imaged presynaptic and postsynaptic activity in the tuft dendrites of L5 ET neurons in premotor cortex as mice performed a cued directional licking task. We defined synapses as belonging to different networks based on a large-scale feature of cortical network topology: how strongly their activity was coupled to the net activity of the population (32). In local cortical networks, a subset of neurons belong to a strongly coupled rich-club with dense intrinsic connectivity (33–37), which may play a unique role in coordinating network dynamics (35, 38). For the first time, we identified a similar topology in the activity of the diverse synaptic inputs to tuft dendrites. We found that tuft spikes were embedded in input episodes with short– and long-timescale components that were highly specific to the dendrite that generated a tuft spike. Tuft spikes were particularly associated with the activity of inputs belonging to the strongly-coupled network or a newly discovered nonpositive-coupled network, networks we found encoded the preparation-to-action transition and the task-outcome, respectively. Together, our results suggest that tuft spikes are associated with activation of distinct subnetworks, which are themselves embedded within a larger-scale cortical network topology.

## Materials and Methods

### Mice and surgeries

All animal procedures were carried out in accordance with protocols approved by the University of Minnesota Animal Care and Use Committee. Twenty adult C57BL/6J mice (Jackson Laboratory; 000664) were used for all experiments (11 male and 9 female). 19 mice were task trained, and 1 mouse was used for imaging during spontaneous behavior. Buprenorphine (0.1 mg/kg, IP injection) and Rimadyl (10 mg/kg, Subcutaneous) were injected 30 minutes prior to surgery for analgesia. Surgical procedures were performed under isoflurane anesthesia (3% for induction, 1–1.5% during surgery). A burr hole was drilled above the left ventromedial thalamus (VM) (coordinates: − 1.5 mm posterior, 0.9 mm lateral relative to bregma). A retrograde adeno-associated viral vector encoding Cre recombinase (pENN-AAVrg-hSyn-Cre-WPRE-hGH 105553; Addgene) was injected into the VM thalamus. The virus was diluted to a final concentration of 1.2 *×* 10^12^ vg/mL, and 200 nL was delivered at a depth of 4000 µm. A 3 mm diameter craniotomy was made above the left hemisphere of ALM (2.5 mm anterior and 1.5 mm lateral to bregma). Cre-dependent adeno-associated viral vectors (pAAV9-hSyn-FLEX-iGluSnFR3-v857-SGZ 175182, pAAV1-Syn-Flex-NES-jRGECO1a-WPRE-SV40 100853; Addgene) were injected into ALM. 2 mice (*n* = 12 sessions) were only injected with iGluSnFR3 and were included in some analyses. (Three 100 nL injections were delivered in a triangular pattern, with injection sites spaced approximately 150 µm apart, at a depth of 700 µm. Viral vectors were diluted to a final concentration of 1.2 *×* 10^13^ vg/mL. A window (triple #1 coverglass 2.5/2.5/3.5 mm diameter; Potomac Photonics, Baltimore, MD) was fixed to the skull using dental adhesive (C and B Metabond; Parkell, Edgewood, NY). A metal bar for head fixation was implanted posterior to the window with a metal loop surrounding the window. Expression was monitored for at least 2.5 weeks before imaging. Mice were gradually water restricted and then received water according to task performance (minimum: 0.6 mL daily) or 1 mL on days without training (39).

### Auditory decision task

Head-restrained mice were trained to perform an auditory-cued directional licking task (Fig. 1A; 39, 40). The behavioral apparatus was controlled using BPOD (Sanworks). Each trial started with the sample period, during which one of two auditory cues (3 kHz or 18 kHz; 5 tones at 4 Hz) was presented. The sample period was followed by a delay epoch of 1.25 s. Following the delay epoch, an auditory “GO” cue (carrier frequency 6 kHz with 360 Hz frequency modulation, 100 ms duration) was presented, which instructed mice to lick one of two water ports centered just below the mouth (2 mm between ports). Licking was detected with a custom electrical lick detector. Licking the correct water port within 2 s (as instructed by the sample cue) dispensed a small amount of water (3 *µ*L) from that port, whereas licking the incorrect port triggered a 15 s timeout period. Licking during the sample or delay period reset the corresponding period timer. Daily sessions generally lasted 1 to 1.5 hours, with 220 to 300 randomized trials. Mice required 3–6 weeks to achieve performance greater than 70%. Imaging sessions began after mice maintained performance above 70% for at least 3 consecutive days.

**Fig. 1.**
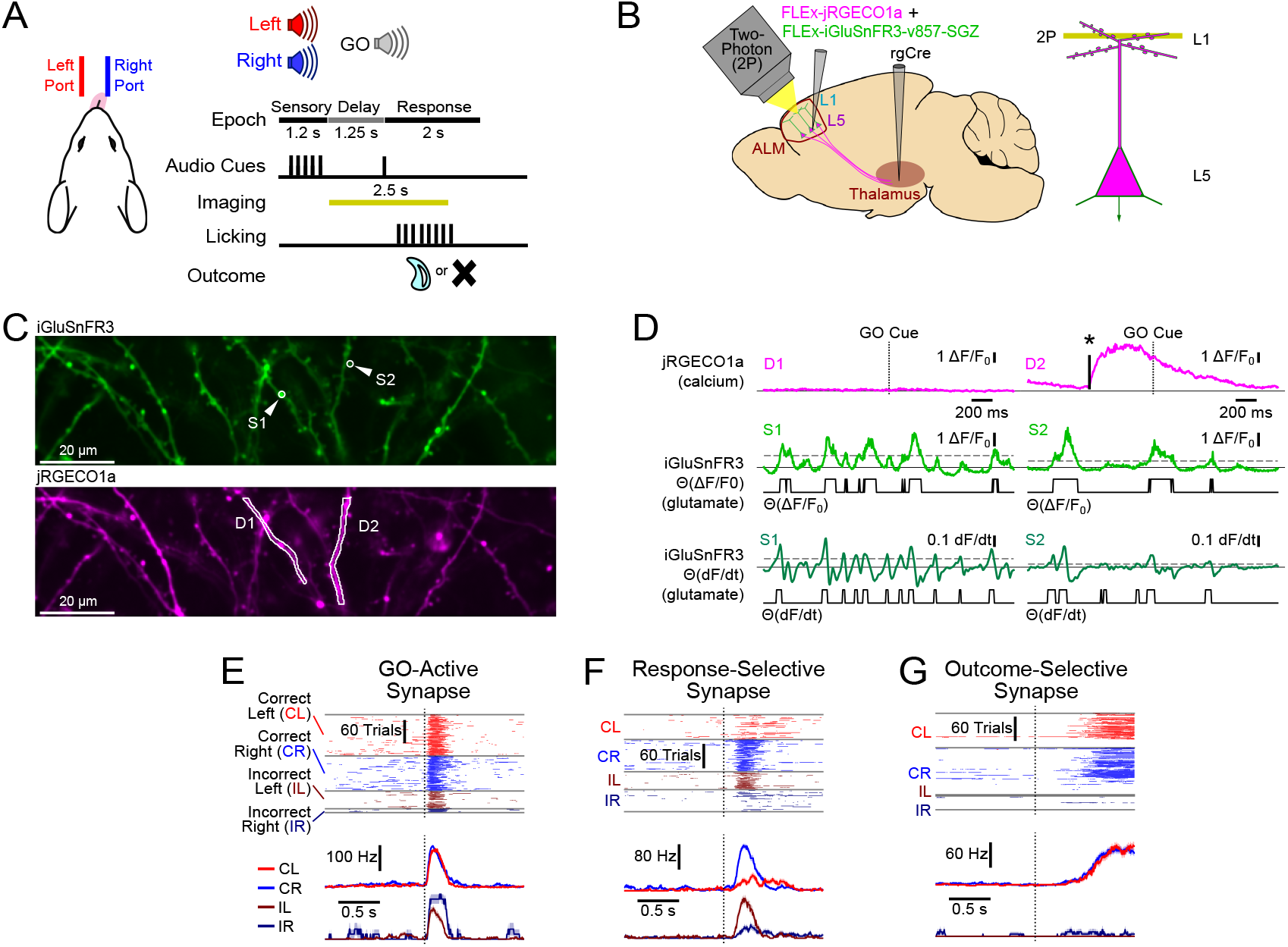
Simultaneous imaging of synaptic input and dendritic events during motor behavior. ***A***, Illustration of cued directional licking task. Yellow line: period of 2P imaging. ***B***, Illustration of labeling and imaging strategy. ***C***, Example FOV showing simultaneously recorded iGluSnFR3 (green; top) and jRGECO1a (magenta; bottom) expression along with synaptic ROIs (S1, S2) and dendritic ROIs (D1, D2). ***D***, Activity for ROIs in (C) for one example trial. Top: jRGECO Δ*F/F*_0_ timecourses of dendrite segments. *: inferred tuft spike onset. Middle: iGluSnFR3 Δ*F/F*_0_ timecourses of synapses (green). Horizontal dashed line shows threshold for Θ(Δ*F/F*_0_) (black). Bottom: iGluSnFR3 *dF/dt* timecourses (dark green). Horizontal dashed line shows threshold for Θ(*dF/dt*) (black). ***E***, Example “GO” active synapse. Top: Θ(Δ*F/F*_0_) traces for individual trials for each trial type. Bottom: trial-averaged timecourses for each trial condition. Error shading: trial-level bootstrap SEM. ***F***, Same as (E), except a “response” selective synapse. ***G***, Same as (E), except an “outcome” selective synapse.

### *In vivo* imaging

Imaging experiments were conducted using a custom-built two-photon microscope utilizing a 25 *×* water-immersion objective (NA 1.05, Olympus XLPLN25XWMP2) and a tunable femtosecond laser (1000 nm; 100 fs; 80 MHz; InsightX3, Spectra Physics). Lateral scanning was performed by a resonant mirror (CRS 12 kHz, Novanta Photonics) conjugated to a set of galvanometer mirrors (6215H 5mm, Novanta Photonics). For simultaneous two-color imaging (dichroic: T565lpxr, Chroma), iGluSnFR3 emission photons (FF01-531/46, Semrock) were collected using a photomultiplier tube (PMT; PMT2101, Thorlabs) and jRGECO1a emission photons (FF01-625/90, Semrock) were collected with a multi-pixel photon counter (MPPC; C14455-0276, Hamamatsu). Frames were acquired at ∼175 Hz over a 140 µm × 35 µm imaging area, digitized (NI-5771/NI-7975R/PXIe-1092, National Instruments), and recorded with ScanImage (MBF Biosciences). For spontaneous activity imaging (Fig. S3E), two sessions were collected at 325 Hz with a 140 µm *×* 17.5 µm new field of view (FOV), and a third session was collected at ∼90 Hz with a 280 µm *×* 140 µm FOV. Imaging was restricted to apical dendrites located less than 0.1 mm below the pial surface. To mitigate rapid photobleaching, imaging was limited to a 2.5 s window per trial, beginning at the start of the delay period and ending 1.25 s into the response period. A new FOV was imaged each day.

### Rigid registration

Image registration methods were as previously described (41). In brief, all image series were subjected to an initial rigid registration (phase cross-correlation; no upsampling; python scikit-image). Offsets were estimated from spatially filtered (Gaussian *σ* ≤ 1 µm) frames, adaptively filtered to suppress registration noise, and then used to transform unfiltered frames.

### Intra-trial bleach correction

To correct for the fast trial-by-trial photobleaching component (Fig. S1A,B), the whole-frame fluorescence time series of each emission channel was averaged across trials. A double-exponential decay was then fit to the mean time series. For some sessions, frames (0-0.4 s) after the GO cue were excluded from the fit to mitigate influence from large transients of activity that occurred at that time. The fit was then reciprocally normalized by the last frame in the fit as:

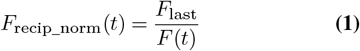

where *t* denotes time (or frame index), *F* (*t*) is the fitted double-exponential photobleaching curve evaluated at time *t, F*_last_ is the value of the fitted photobleaching curve at the final frame included in the fit, and *F*_recip_norm_(*t*) is the reciprocal normalization factor used to correct fluorescence signals for photobleaching. Then it was multiplied element-wise to the activity traces of each trial (Fig. S1B).

### Denoising

Following rigid registration, image series were denoised using DeepInterpolation (42), modified to minimize L2 (rather than L1) loss. Denoising was performed separately on glutamate and Ca^2+^ activity channels. For interpolation of jRGECO1a frames, the 3 preceding and 1 following frames were provided for training and inference. For interpolation of iGluSnFR3 frames, 6 mean projections of frames collected from non-overlapping exponential (base 1.26; exponents: 0–5) frame windows both preceding and following the interpolated frame were provided for training and inference. Following denoising, the first and last 12 frames from each trial were excluded to remove artifacts of interpolation near the edges of the 2.5 s imaging windows.

### Non-rigid registration

Small within-session instabilities (*e*.*g*., resonant mirror temperature changes, vessel dilation) resulted in image warp that was significant for dendritic imaging. Image warp was corrected as previously described (41). In brief, warp calculations were performed on contrast-enhanced frames using symmetric diffeomorphic registration (python DIPY; 43, 44), adjusting the spatial scale of the warp correction on a session-by-session basis. To mitigate the noise of frame-to-frame warp calculations, *x*– and *y*-warp fields were adaptively filtered in time.

### Activity extraction

Dendrite regions of interest (ROIs) were manually segmented from the mean projection image of each session. Active synapses were manually segmented from the variance projection image of each session and assigned to dendrite segments based on the mean projection image. Only dendrites and synapses within regions of the imaging FOV where labeling was sufficiently sparse to permit clear assignment of synapses to dendrite segments were segmented. Mean fluorescence within each ROI was divided by rolling baseline (*F*_0_; averaging window: *±* 7 trials; synapse activity exclusion: z-score *>* 2; dendrite activity exclusion: z-score *>* 3) and subtracted by 1 to obtain Δ*F/F*_0_. Synapse SNR was estimated as the value of the 99th percentile of positive Δ*F/F*_0_ deflections divided by 99th percentile of the absolute value of negative Δ*F/F*_0_ deflections. Synapses with SNR *<* 1.5 were excluded.

Two complementary transformations of iGluSnFR3 activity were used to assess synaptic activity. One was derived from the conventional Δ*F/F*_0_ and the other was derived from the smoothed *dF/dt* (Gaussian filter *σ* = 2 with a radius of 4 frames). Both measures were binarized by a shifted Heaviside step function Θ(*x* − *x*_*neg*%_), where *x* is the continuous measure (Δ*F/F*_0_ or *dF/dt*) and *x*_*neg*%_ is the 99th (for Δ*F/F*_0_) or 92^nd^ percentile (for *dF/dt*) of the absolute value of negative deflections *x*. To obtain the trial-specific activity (*TSA*) of a synapse, the trial-averaged activity for each trial type (CL, CR, IR, IL) was subtracted from activity on corresponding trials (Fig. 2G).

**Fig. 2.**
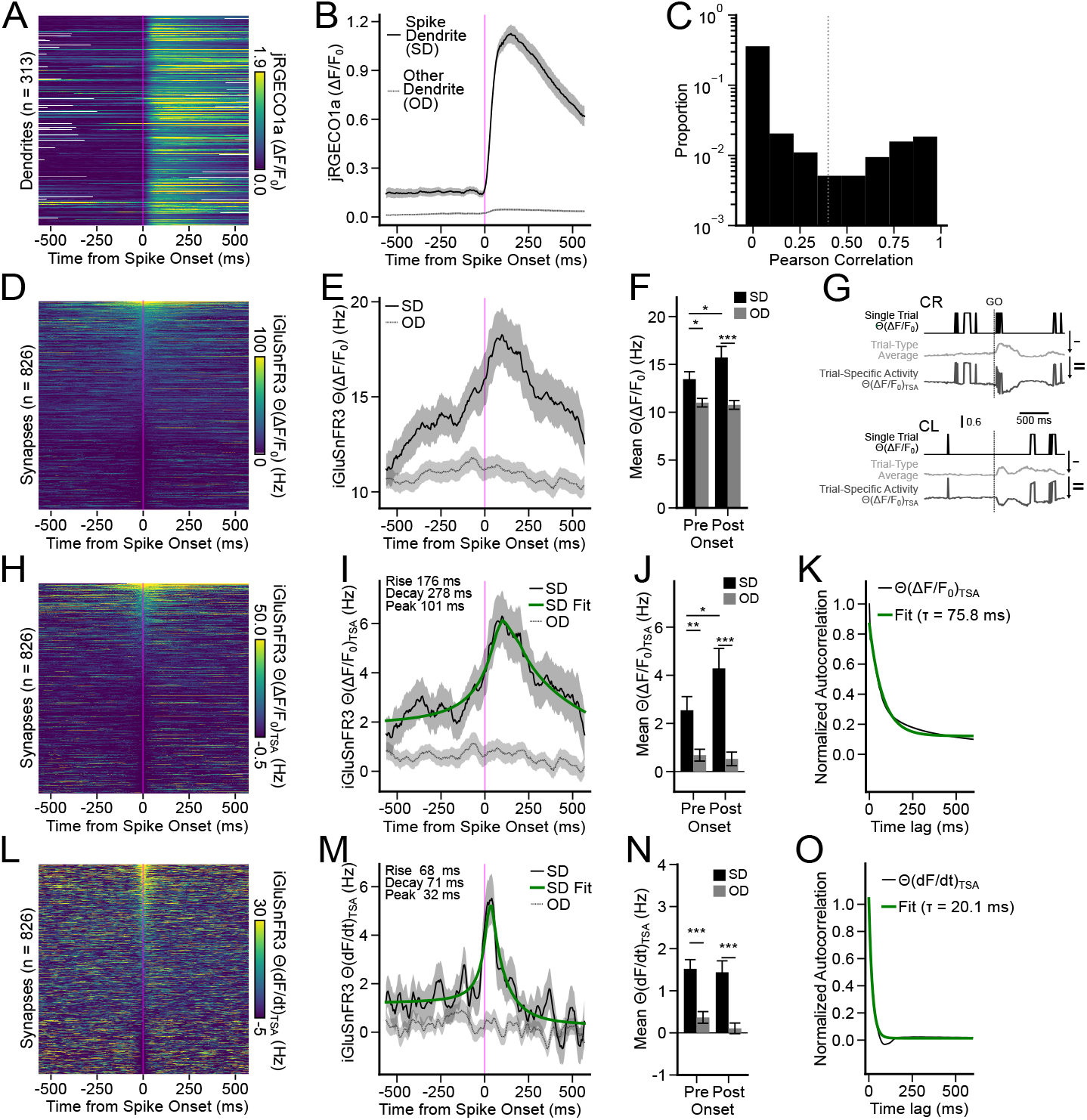
Tuft spikes are associated with temporally extended, dendrite-specific increases in input. ***A***, Trial-averaged spike dendrite (SD) Ca^2+^ transients aligned to inferred spike onset for individual dendrites. ***B***, Mean population SD Ca^2+^ transient for dendrites in (A) (solid black; *n* = 313 dendrites) or other dendrite (OD) Ca^2+^ transient (dashed gray; *n* = 509 dendrites). ***C***, Pairwise dendritic Ca^2+^ correlations (*n* = 5078 pairs) with gray line at 0.4 correlation. ***D***, Trial averaged SD Θ(Δ*F/F*_0_) of individual synapses. ***E***, Population mean Θ(Δ*F/F*_0_) of SD synaptic activity (black solid line; *n* = 826 synapses) and OD synaptic activity (dashed gray; *n* = 1922 synapses) aligned to tuft spikes. Green line: fit of SD to exponential rise and decay (*R*^2^ = 0.92). ***F***, Mean of pre-onset (–575–0 ms) and post-onset (0–575 ms) activity in (E). ***G***, Illustration of the calculation of trial-specific activity (TSA, see Methods). ***H-J***, Same as (D-F), but for Θ(Δ*F/F*_0_)_*T SA*_. ***K***, Mean normalized autocorrelation of Θ(Δ*F/F*_0_)_*T SA*_ across synapses (black; *n* = 2072 synapses) and exponential decay fit (green; *R*^2^ = 0.98). ***L-O***, Same as (H-J) for Θ(*dF/dt*)_*TSA*_ activity. Fit in (M): *R*^2^ = 0.81. Fit in (O): *R*^2^ = 0.98. Error shading and error bars are session-level hierarchical bootstrap SEM (see statistics in Table S1).

Tuft spikes were detected in the jRGECO1a dendrite segment Δ*F/F*_0_ timecourses using a custom algorithm. Transients above the 99.9^th^ percentile of negative Δ*F/F*_0_ deflections for 15 consecutive frames were identified as potential spikes. Subsequent to the first transient identified in a trial, transients were only considered as potential spikes if activity dropped back below threshold for at least 25 frames prior to the transient. To identify the onset time for each potential spike, the derivative of a 13-frame moving average of Δ*F/F*_0_ was convolved with an odd-symmetric slope kernel [2, 2, 2, 1, 1, 1, 0, −1, − 1, − 1, − 2, −2, −2] and then smoothed again with a 13-frame moving average. Within the resulting timecourse, the maximum value within the 30 frames preceding the first frame of the transient was defined as the onset time. Only spikes with a relative Δ*F/F*_0_ increase of at least 0.3, measured as the difference between the onset frame Δ*F/F*_0_ and the average Δ*F/F*_0_ across frames 12–25 after onset, were retained.

### Pairwise correlation and autocorrelation analyses

Pairwise correlation of dendrite segment Ca^2+^ activity was determined using floored Δ*F/F*_0_ timecourses (Fig. 2C). Values below the 99.9^th^ percentile of the absolute value of negative Δ*F/F*_0_ deflections were zeroed, and values above were left unchanged. Floored traces were used to ensure that small motion artifacts did not dominate the correlation estimate, as most dendrites produced few transients per session (Fig. S1C). Pairwise correlation of synaptic activity was based on session-wide Θ(*dF/dt*)_*TSA*_ timecourses for all analyses (e.g. Fig. 3D). Synapse pairs within 2.5 µm in Euclidean distance were excluded to avoid contamination of input signals by potential glutamate spillover. Synapse pairs with a correlation *>* 0.5 were also excluded to remove synapses potentially originating from the same axon. In all cases, Pearson’s correlation coefficient was used as the correlation metric.

**Fig. 3.**
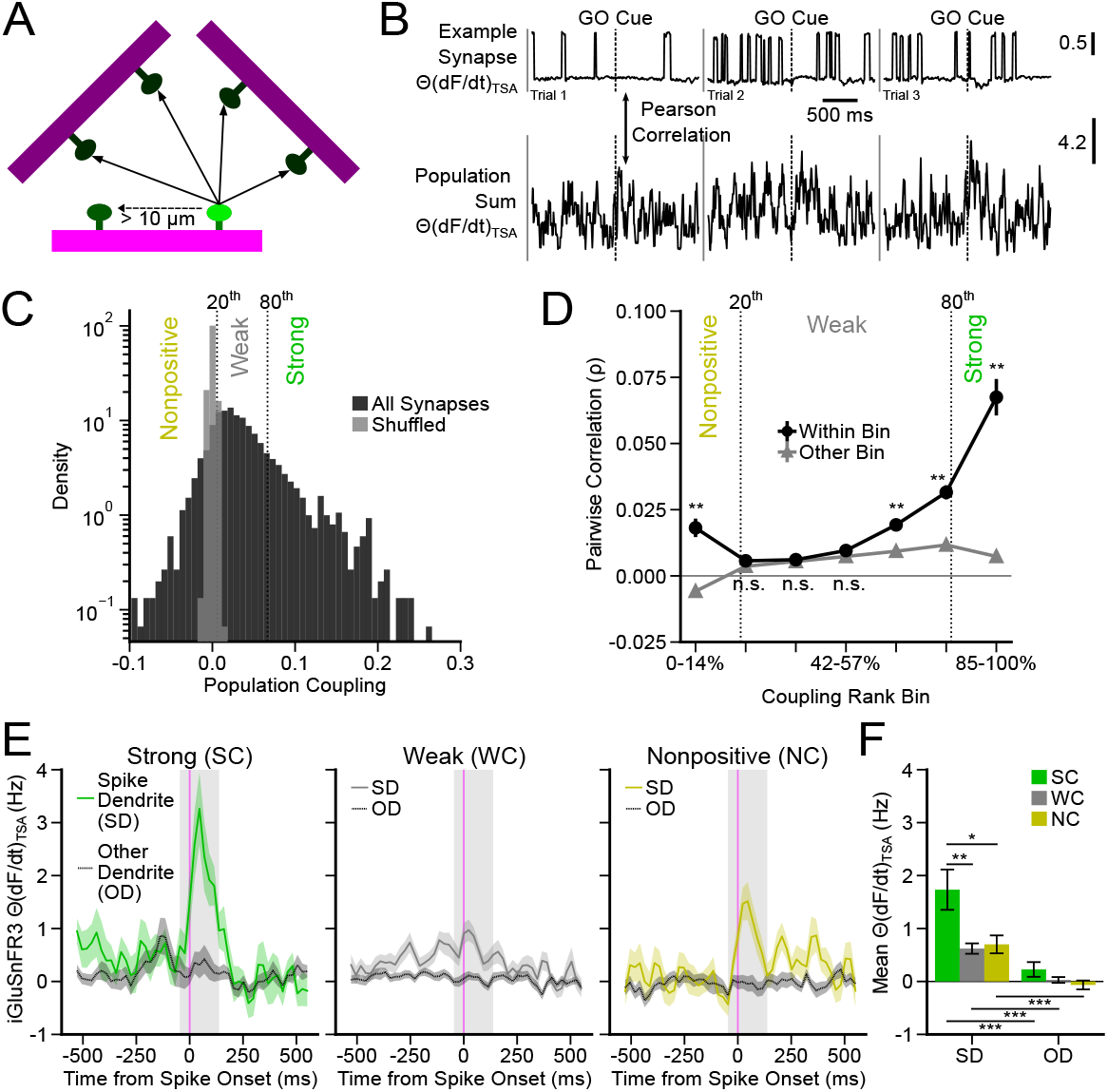
Strongly coupled synapses are particularly active at tuft spike onset. ***A***, Illustration of the synapse population used to calculate the coupling of one synapse (bright green). ***B***, Example snippet of Θ(*dF/dt*)_*TSA*_ from a synapse and corresponding synapse population sum Θ(*dF/dt*)_*TSA*_. ***C***, Distribution of population coupling values across all synapses (black; *n* = 2072 synapses) and distribution after shuffling time (gray). ***D***, Mean pairwise correlation (*ρ*) calculated from Θ(*dF/dt*)_*TSA*_ of synapse pairs within the same bin (black circles; *n* = 9561 synapse pairs) or with synapses in other bins (gray triangles; *n* = 71212 synapse pairs) binned by population coupling rank (statistics in Table S2). ***E***, Θ(*dF/dt*)_*TSA*_ synaptic SD activity (solid lines) and OD activity (dashed lines) aligned to tuft spikes for synapses grouped by coupling strength (*n*: *SC*_*SD*_ = 144; *SC*_*OD*_ = 382, *WC*_*SD*_ = 522; *WC*_*OD*_ = 1156; *NC*_*SD*_ = 160; *NC*_*OD*_ = 384). ***F***, Mean activity in gray windows (–50–150 ms from spike onset) shown in (E). Significance tests: two-tailed hierarchical difference test (statistics in Table S3). Error shading and error bars: session-level hierarchical bootstrap SEM. All panels only included data with dual color imaging.

The autocorrelation of individual synapses was calculated using Pearson’s correlation coefficient at increasing time lags of Θ(*dF/dt*)_*TSA*_ or Θ(Δ*F/F*_0_)_*TSA*_, as noted (Fig. 2K,O). The autocorrelation decay constant was calculated from a single-exponential fit (with intercept) to the mean of the autocorrelation traces of all synapses. It was estimated by iteratively fitting the model to resampled group-level means (see Statistics section for details on bootstrapping procedures). The zero-lag autocorrelation value was excluded from this fit to mitigate distortion of the estimate by photon shot noise.

### Characterization of synaptic activity associated with tuft spikes

Synaptic activity was aligned to the inferred onset of tuft spikes and averaged for each synapse before calculating group-level summary statistics. Only synapses on dendrite segments where at least 4 tuft spikes were detected across the imaging session were included in this analysis. For each spike, synapses located on the dendrite in which the tuft spike was detected were considered part of the spike dendrite group (SD). The other dendrite group (OD) consisted of simultaneously imaged synapses detected on other (defined by session-wide calcium activity correlation *<* 0.4) dendrite segments (Fig. 2C). Putative multi-branch tuft spikes were defined as events where another dendritic segment in the FOV had a session-wide calcium activity correlation *>* 0.4 and a same-trial correlation *>* 0.7 with the spike dendrite (Fig. S1J,K).

Synaptic dynamics associated with tuft spikes were quantified using a composite model comprising a single-exponential rise and a single-exponential decay with a joint intercept. The fit of the composite model was estimated by iteratively fitting the model to resampled group-level means (see Statistics section for details on bootstrapping procedures). For each bootstrap sample, the transition point between the rise and decay phases was defined as the time of the maximum value within 0–400 ms of tuft spike onset.

### Population coupling

Population coupling estimation used Θ(*dF/dt*)_*TSA*_ timecourses, with the exception of estimation from spontaneous activity (Fig. S3E) which used Δ*F/F*_0_. Population coupling was quantified by computing the Pearson correlation coefficient between a synapse and the summed activity of all other simultaneously imaged synapses, excluding those located within 10 µm on the same branch (Fig. 3A,B). Synapses located nearby were excluded to avoid functional sampling biases. For the calculation of population coupling rank, all synapses across all sessions were pooled. Synapses were divided into three groups based on population coupling rank: nonpositive (bottom 20%), weak (middle 60%), and strong (top 20%). The shorthand identifier “nonpositive” was adopted since this group contained synapses with a population coupling that was either negative or statistically indistinguishable from zero.

### Activity modes and coding directions

The GO mode is a dimension in the space of synaptic activity that best captures the lick-direction-invariant change in activity pattern following the GO cue, which contributes to transitioning population activity from a motor-preparatory to an action-ready state (45). The Response Direction is a direction in activity space that maximally distinguishes between activity on CR and CL trials (45, 46). The Outcome Direction is a direction in activity space that maximally distinguishes between activity on correct and incorrect trials (47, 48). A weight for each synapse for each mode or direction was calculated as follows: **GO mode:**

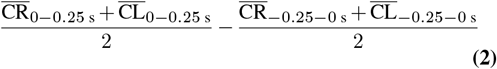

**Response Direction:**

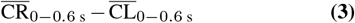

**Outcome Direction:**

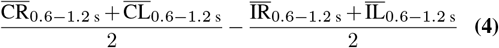

where 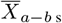 denotes the trial-averaged synaptic activity of trial-type *X* over the time interval from *a* to *b* seconds relative to the GO cue. The GO mode was orthogonalized to the Response Direction using a Gram-Schmidt process to isolate condition-invariant activity. The Outcome Direction was orthogonalized to the GO mode and then to the Response Direction using a Gram-Schmidt process.

### Analysis of synapse clustering by function

Pairwise noise correlation as a function of distance analysis was calculated using Θ(*dF/dt*)_*TSA*_ timecourses (Fig. S2A). Signal correlation was calculated using trial-averaged Θ(*dF/dt*) time courses of each trial-type concatenated together (Fig. S2B). Distance was measured along the centerline of dendritic segments for pairs located on the same dendrite. Euclidean distance was used for synapses located on different dendrites. The spatial structure of pairwise correlations was quantified by fitting a single-exponential decay model. The model was iteratively fit to resampled group-level means (see Statistics section for details on bootstrapping procedures).

To determine whether strongly coupled synapses were preferentially clustered on the same dendritic branch (Fig. S2C), for each strongly coupled synapse, we evaluated pairwise relationships with other synapses located either on the same dendrite or on different dendrites. Pairs in which both synapses were classified as strong couplers were assigned a value of 1, whereas pairs containing a weak or nonpositive coupler were assigned a value of 0. These values were then averaged separately across same-dendrite and different-dendrite pairs for each reference strong coupler synapse. To avoid biases arising from unequal sampling, we balanced the number of pairs of same-dendrite and different-dendrite for each reference synapse prior to averaging.

### Secondary analysis of electrophysiology dataset

We reanalyzed publicly available electrophysiology DANDI data that measured responses of single neurons in mouse ALM during a similar behavioral task (Fig. S4; 49, 50). Detailed methods are included with the data. In brief, mice engaged in a similar delayed directional licking task, except that the sensory cue was tactile. Neural activity in ALM cortex was recorded with NeuroNexus silicon probes and spike sorted. Our analysis included eight Sim1_KJ18-Cre mice and three Tlx_PL56-Cre mice from this study. Only high-quality units, as labeled in the original dataset, were included. To obtain more reliable coupling estimates, we also included only sessions that contained at least 18 units with a session-wide firing rate *>* 0.5 Hz. Trials with photostimulation were excluded. Spike trains were downsampled by counting spikes in 5 ms bins and smoothed with a Gaussian filter (*σ* = 2 bins; 20 ms radius).

Population coupling for neurons was computed using procedures closely matching those described for the synaptic data (Fig. S4B). Trial-specific activity was calculated as described for synaptic imaging data. The population coupling of each neuron was estimated session-wide from activity recorded between the tactile stimulus onset and 2 s after GO signal.

### Brain motion exclusion analysis

For some control analyses (Fig. S1G,H; Fig. S3D), we blanked inferred large motion periods to establish that dynamics we observed were not explained by motion artifacts. Large motion periods were inferred from the *x* and *y* registration corrections since animal movements are expected to be correlated in multiple directions. For each frame, the absolute *x* and *y* registration corrections were summed. To account for slow drift, a rolling movement metric (Δ*M/M*_0_) was computed by averaging motion from the current trial together with the seven preceding and seven subsequent trials. For edge trials, the maximum number of available neighboring trials in the constrained direction was used. The resulting Δ*M/M*_0_ trace was smoothed with a Gaussian filter (*σ* = 2, radius = 5 frames) prior to computing the first derivative. The absolute value of this derivative served as the final motion estimate. Motion traces from all experimental sessions were concatenated to establish a uniform threshold (*>* 85 ^th^ percentile) based on the highest-motion periods across the full dataset (Fig. S1G). This prevented over-penalization of low-motion sessions and under-penalization of high-motion sessions.

### Statistics

Group-level summary statistics were computed using nonparametric bootstrap procedures to avoid assumptions of normality. Group means, error bars, and mean differences were estimated using hierarchical bootstrapping. For each bootstrap resample, imaging sessions were sampled with replacement until the original number of observations was drawn. Observations were subsequently averaged at the lower hierarchical level specified in the corresponding figure or text. Unless otherwise noted, 10,000 valid (*i*.*e*. non-NaN) resamples were conducted for all analyses. For activity timecourses, the mean of each resample was lightly smoothed using a Gaussian filter (*σ* = 2, radius = 4 frames) or binned by averaging every four frames (for coupling group timecourses).

For primary pairwise coupling and correlation analyses (Fig. 3D), synapse pairs were generated by resampling individual synapses 100 *×* without replacement (drawing unique, non-overlapping subsets so each synapse contributed to at most one pair within a resample). These estimates were averaged to obtain a single value per bootstrap sample and this procedure was repeated for 1,000 bootstrap iterations that resampled imaging sessions with replacement.

For mean difference tests, group comparisons were performed directly on the bootstrap difference distributions. For each comparison between groups *i* and *j*, bootstrap means were paired by iteration and the difference distribution was computed as *d*_*k*_ = *a*_*k*_ − *b*_*k*_, where *a*_*k*_ and *b*_*k*_ denote the bootstrap estimates from iteration *k* for the two groups. To avoid *p*-values of exactly zero due to finite bootstrap sampling, each tail probability was computed with the add-one correction (51) as (*c* + 1)*/*(*n* + 1), where *c* is the number of bootstrap differences in the relevant tail and *n* is the number of valid (*i*.*e*. non-NaN) bootstrap differences. One-tailed tests evaluated the *a priori* hypothesis that group *i* exceeded group *j* by calculating the proportion of bootstrap differences less than or equal to zero, *p*(*d*≤ 0). Two-tailed tests were computed as 2 ×min(*p*(*d*≤ 0), *p*(*d*≥ 0)), with values capped at 1.0. When multiple pairwise comparisons were performed, raw bootstrap *p*-values were corrected for multiple testing using the Holm procedure, and corrected *p*-values below *α* = 0.05 were considered statistically significant. For relevant figures, asterisks indicate statistical significance based on the Holm-corrected *p*-value (*∗ p <* 0.05, *∗∗ p <* 0.01, and *∗∗∗ p <* 0.001).

## Results

### Tuft spikes are accompanied by dendrite-specific, temporally extended input episodes

To measure relationships between excitatory input, tuft spikes, and cortical dynamics, we imaged presynaptic and postsynaptic activity in the anterior lateral motor (ALM) cortex of mice performing a delayed directional licking task (Fig. 1A, 39). During this behavior, task-driven activity patterns in ALM cortex exhibit well-described periods of both extended stability and rapid transition (52, 53), making it an especially suitable region to evaluate the relationship between tuft spikes and input dynamics. To precisely measure the timing of presynaptic glutamatergic input, we took advantage of a recently developed genetically encoded glutamate sensor with an affinity well-suited to report fast synaptic glutamate transients (iGluSnFR3, 54, 55). To selectively label L5B ET neurons, we injected a retrograde AAV-Cre virus into motor thalamus, as well as Cre-dependent AAVs encoding iGluSnFR3 (a green indicator) and jRGECO1a (a red calcium indicator) into ALM (Fig. 1B). We focused on L5B ET neurons because they have prominent apical tuft dendrites (56, 57) and because restricting labeling to these neurons ensured that any dendrite-specific synaptic activity we observed did not simply reflect activity for the tuft dendrites of neurons belonging to one sublamina over another.

In mice already expert at the task, we conducted daily highspeed (175 Hz frame rate) imaging of the dendrites in L1 (Fig. 1B,C) during the performance of hundreds of behavioral trials (median = 235; range = 162–302), selecting a new fieldof-view (FOV) for imaging each session (N=19 mice, 64 FOV, 611 dendrite segments, 2591 synapses). 2 mice (*n* = 12 FOVs) only expressed iGluSnFR3 and were included in some analyses. Because iGluSnFR3 undergoes rapid photobleaching (54), we restricted imaging to a 2.5 s window spanning the beginning of the delay epoch to 1.25 s after the GO cue (Fig. 1A). Image series were corrected for brain motion, denoised, and manually segmented (Fig. 1C; see Methods). Extracted fluorescence timecourses were then corrected for the effects of bleaching (Fig. S1A,B; see Methods). We analyzed synaptic signals using two different binarized (noise-shifted Heaviside function Θ, see Methods) transformations of the iGluSnFR3 fluorescence. The first measure derived from the conventional measurement of change from baseline (Θ(Δ*F/F*_0_)) that presumably reflected cumulative glutamate release as convolved by iGluSnFR3 dynamics. The second measure derived from the positive derivative of fluorescence (Θ(*dF/dt*)) that presumably reflected rapid increases in glutamate release after periods of relative quiescence (Fig. 1D). The trial-averaged activity of individual synapses (Fig. 1E-G) exhibited selectivity for different task-variables similar to those previously reported for neurons in ALM cortex (47, 53), including synapses consistently driven by the GO cue (Fig. 1E), selective for lick direction (Fig. 1F), and selective for task-outcome (Fig. 1G).

To align synaptic activity with the occurrence of tuft spikes, we inferred the timing of tuft spikes from the onset of transient increases in the jRGECO1a fluorescence within the dendritic shaft (Fig. 1D; Fig. 2A,B; see Methods). For the purposes of this study, the term “tuft spike” refers to any regenerative activity within the tuft dendrites, and is therefore agnostic to the spatial extent, engaged conductances, or genesis of the activity. Dendrite segments were much larger than individual synapses (median = 18 µm; range = 4-105 µm) and our inference algorithm only considered large jRGECO1a transients (*>*30% Δ*F/F*_0_) as potential spikes, so it is unlikely that the activity of individual synapses or subthreshold cooperativity of a few synapses were mistakenly identified as regenerative events. Consistent with this, tuft spikes were detected at rates ≈100 × lower than synaptic activity (Fig. S1C), also suggesting that most bAPs were not detected as tuft spikes. Dendrite segments putatively originating from two different neurons were identified by a pairwise calcium session correlation of less than 0.4. This threshold was the approximate antimode within the bimodal distribution of correlations between segments (Fig. 2C), and previous work has found that even the most independently active compartments of the tuft dendrite exhibit average calcium signal correlations greater than 0.4 (23).

The timecourse of mean synaptic activity (Θ(Δ*F/F*_0_)) aligned to tuft spikes was highly variable across individual synapses (Fig. 2D), however, averaging across all synapses revealed a clear peri-spike elevation in synaptic activity (Fig. 2E). Compared to simultaneously recorded dendrites of putative other neurons – henceforth simply referred to as “other” dendrites (OD) – mean synaptic activity on the spiking dendrite (SD) was significantly elevated both before (*SD* = 13.5 *±* 0.7; *OD* = 11.0 *±* 0.4 Hz; *p* = 0.002) and after (*SD* = 16 ±1.0; *OD* = 10.8 ±0.4 Hz; *p* = 6.0× 10^−4^) the spike (Fig. 2F; Table S1). These dynamics could simply reflect the relative timing of trial-aligned activity in dendrites and their respective synaptic inputs. To determine if trial-to-trial fluctuations in synaptic activity were associated with tuft spikes, the mean trial-aligned activity of each synapse was subtracted from its timecourses before alignment with tuft spikes (Fig. 2G). Spike-associated increases in this trial-specific activity, Θ(Δ*F/F*_0_)_*TSA*_, were at least 5 × larger on the spiking dendrite than others, both before (*SD* = 2.6 ±0.5; *OD* = 0.7± 0.2 Hz; *p* = 6.0 ×10^−4^) and after (*SD* = 4.3 ±0.7 Hz; *OD* = 0.5± 0.3 Hz; *p* = 6.0× 10^−4^) the spike (Fig. 2H-J; Table S1). Restricting analysis to spikes with larger peak Δ*F/F*_0_ (Fig. S1D-F; Table S6) or excluding frames with large lateral brain motion (Fig. S1G-I; Table S6) did not significantly change the spike-associated synaptic activity timecourses. Mean spike-associated Θ(Δ*F/F*_0_)_*TSA*_ dynamics were well-fitted by an exponential rise (176± 79 ms, half-width at half-max; HWHM) and decay (278 ± 79 ms, HWHM) with a peak that lagged (101±38 ms) the tuft spike (Fig. 2I). These dynamics were substantially longer than the reported iGluSnFR3 kinetics (rise constant *<*2 ms; decay constant = 29 ms; 55) and the mean autocorrelation time constant of the Θ(Δ*F/F*_0_)_*TSA*_ signal (Fig. 2K), indicating that tuft spikes were indeed associated with a dendrite-specific elevation in glutamatergic input activity that extended hundreds of milliseconds before and after the spike.

Abrupt increases in excitatory input are generally more effective at triggering regenerative activity than gradual increases in input (13, 58, 59). To determine whether abrupt increases in synaptic activity from trial-to-trial were associated with tuft spikes, we calculated trial-specific activity of the binarized positive derivative, Θ(*dF/dt*)_*TSA*_. The spike-associated increase in Θ(*dF/dt*)_*TSA*_ exhibited a more rapid rise (68±49 ms, HWHM) and decay (71 ± 26 ms, HWHM) than the spike-associated increase in Θ(Δ*F/F*_0_)_*TSA*_, with a peak that was closer (32±13 ms) to spike onset (Fig. 2L-O). This increase was specific to the spiking dendrite (Fig. 2N; Table S1), both before (*SD* = 1.5 ± 0.2; *OD* = 0.4 ± 0.1 Hz; *p* = 6.0 ×10^−4^) and after (*SD* = 1.4 ± 0.3; *OD* = 0.1± 0.1 Hz; *p* = 6.0×10^−4^) spike onset. Thus, in addition to being associated with an extended elevation of dendrite-specific input activity, tuft spikes were tightly associated (within ≈ 140 ms) with trial-to-trial abrupt increases in input activity that was highly dendrite-specific.

The high dendrite-specificity of the spike-associated synaptic activity could reflect synchronous activity among adjacent inputs directly driving local dendritic spikes. Indeed, pairwise trial-to-trial and task-driven correlations between synapses did decline as a function of distance (Fig. S2A,B; 19, 60). However, most calcium transients detected in the dendritic shaft of the tuft dendrites extend across large regions of the tuft (19, 22, 23). When we restricted our analysis to dendrites with multiple branches in the FOV and multi-branch calcium transients (see Methods), the synaptic activity associated with these confirmed multi-branch events was still dendrite-specific (Fig. S1J-K; Table S6). This suggests that a significant portion of the activity associated with tuft spikes may be selective for the broader tuft dendrites of particular postsynaptic neurons, rather than solely selective at the level of small dendritic segments.

In summary, tuft spikes were associated with both short-timescale and long-timescale increases in glutamatergic synaptic activity on the spiking tuft, while synaptic activity changed minimally on the tuft dendrites of other neurons belonging to the same L5B subclass.

### Inputs strongly coupled to the population are the most active at tuft spike onset

After establishing that an abrupt increase in input was associated with the precise timing of large-scale tuft spikes, we sought to identify functionally-defined networks that might contribute to synchronizing this input. The activity of some cortical neurons is strongly coupled to the activity of the population (32) and these neurons are densely connected to each other, forming a “rich-club” topology (37). We hypothesized that the activity of highly coupled synapses – synchronized by their dense connectivity – might be particularly associated with the generation of tuft spikes.

We calculated the population coupling of each synapse as the correlation between its Θ(*dF/dt*)_*TSA*_ and the net Θ(*dF/dt*)_*TSA*_ of all other synapses within the FOV, excluding synapses on the same dendrite within 10 µm (Fig. 3A,B) to minimize any sampling bias introduced by distance-dependent pairwise correlations (Fig. S2A,B). Similar to the distribution of coupling among local cortical neurons (32), the coupling of synapses to the L5 ET tuft input population was centered near zero, with a long tail of highly coupled synapses (Fig. 3C). Synapses from all sessions were then pooled and assigned percentile ranks based on coupling strength. This coupling rank estimate was robust to various potential technical artifacts, including synapses per FOV, signal amplitude, and brain motion (Fig. S3A-D). We next asked whether synapses with similar coupling strengths were preferentially coactive. As found in local cortical populations (32), highly coupled synapses were more correlated with each other than with other synapses (Fig. 3D; Table S2). Surprisingly, synapses with the lowest coupling also exhibited a higher correlation among themselves (Fig. 3D; Table S2). These enhanced correlations among the lowest couplers were also found in continuous imaging of spontaneous synaptic activity (Fig. S3E; Table S7) and publicly available silicon probe recordings of ALM activity (Fig. S4; Table S8; 49), indicating that this may be a general feature of cortical activity rather than specific to our task, labeled synaptic population, or recording method. Thus, our data suggest the existence of two large-scale functional networks composed of synapses at opposite extremes of the distribution of population coupling. To pool synapses into groups with respect to this organization, we categorized synapses based on their rank coupling as strongly coupled (*SC*; top 20%), weakly coupled (*WC*; middle 60%), or nonpositive coupled (*NC*; bottom 20%).

Next, we compared mean Θ(*dF/dt*)_*TSA*_ aligned to tuft spike onset across coupling groups (Fig. 3E-F). SC synapses exhibited a higher rate of abrupt increases in activity at tuft spike onset (window: –50 ms to +150 ms) than the other coupling groups (*SC*_*SD*_ = 1.7 ±0.4 Hz; *WC*_*SD*_ = 0.6± 0.1 Hz; *NC*_*SD*_ = 0.7 ±0.2 Hz; *SC*_*SD*_ vs. *WC*_*SD*_: *p* = 0.002; *SC*_*SD*_ vs. *NC*_*SD*_: *p* = 0.02). Interestingly, this spike-associated increase in SC activity was still highly dendrite-specific (*SC*_*OD*_ = 0.2± 0.1 Hz; *SC*_*SD*_ vs. *SC*_*OD*_: *p* = 6 ×10^−4^). This suggests that although SC synapses are relatively strongly correlated with each other (same coupling range = 0.067 ±0.007; different coupling range = 0.007 ±0.002; *p* = 7.0 × 10^−3^; Fig. 3D), at the onset of the tuft spikes the activity of SC synapses is driven along a dendrite-specific dimension in the activity space, and this dimension is distinct from the dimension that best captures their population-wide coactivity. Although there was a weak tendency for synapses of the same coupling group to cluster on the same dendrite segment, it was not statistically significant (mean difference: –0.053; *p* = 0.19; Fig. S2C), suggesting that SC synapses were not strongly targeted to particular dendrite segments.

### Spike-associated activity occurs along distinct dimensions partitioned by task-selectivity and population

#### coupling

After discovering a relationship between the activity of synapses at the onset of tuft spikes and the population coupling of synapses, we sought to determine whether different coupling groups encoded distinct task-variables. Trial-averaged Θ(Δ*F/F*_0_) timecourses revealed clear qualitative differences in the timing of activity between coupler groups (Fig. 4A-F; Fig. S5A). SC synapses were predominantly active immediately following the GO cue (Fig. 4D), whereas NC synapses were predominantly active during a later epoch when mice typically collected reward (Fig. 4F). The timing of WC synapse activity was intermediate to the two other groups (Fig. 4E). Individual synapses of the different coupling groups also had qualitatively distinct selectivity for trial-type (Fig. 4A-C). Importantly, these differences in the average magnitude of early-response activity did not trivially define synaptic coupling, as coupling ranks remained stable after excluding this epoch (Fig. S3F).

**Fig. 4.**
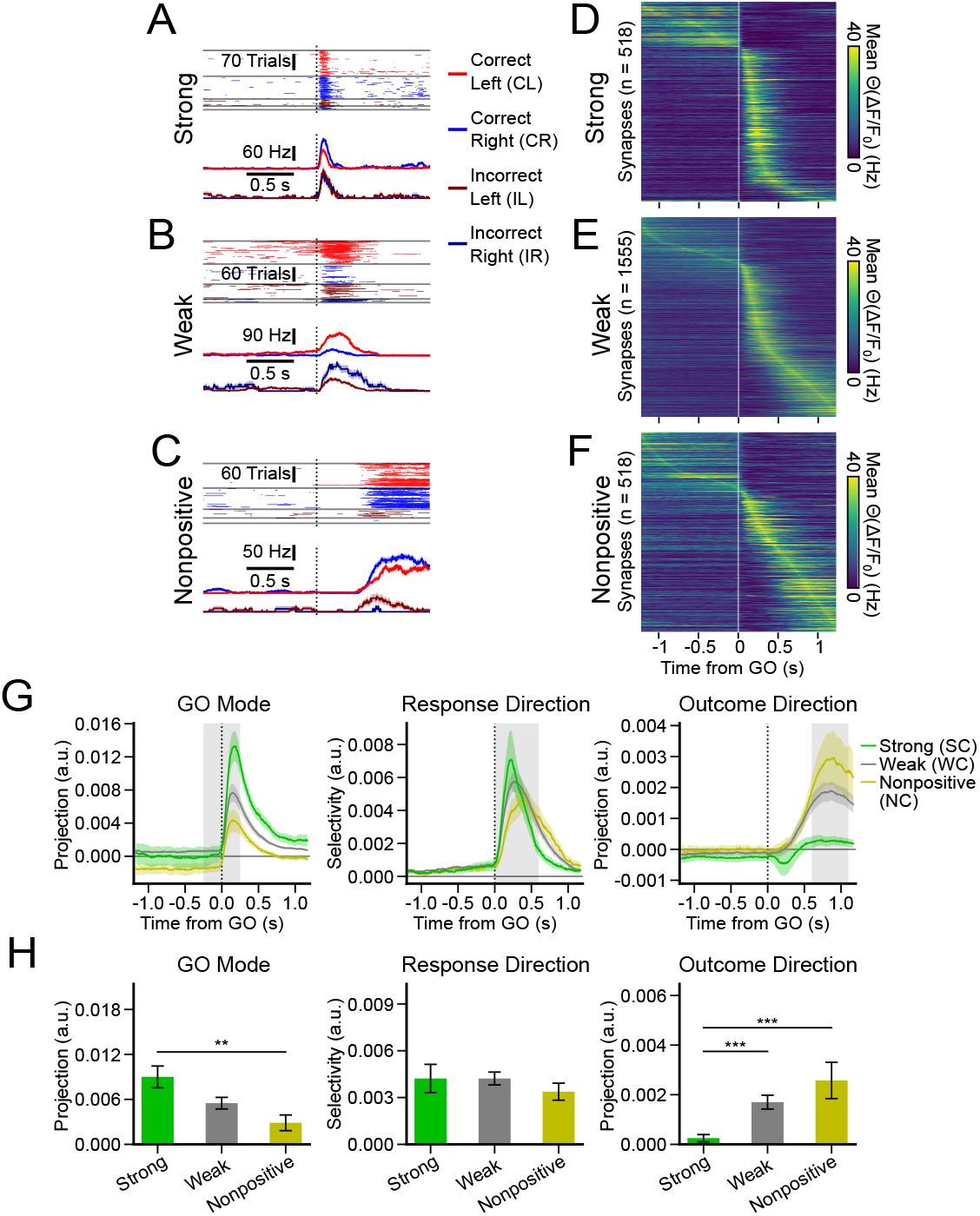
Synapses with different population coupling have distinct task-selectivity. ***A***, Example strong coupling (SC) synapse showing condition-invariant “GO” activity. Top: Θ(Δ*F/F*_0_) activity for individual trials for each trial condition. Bottom: trial averaged time courses for each trial condition. Error shading: bootstrap trial SEM. ***B***, Same as (A) except a WC synapse with lick-direction selective activity. ***C***, Same as (A) except an NC synapse showing positive reward-outcome activity. ***D-F*** Peak-time-sorted heatmaps of trial averaged time courses (Θ(Δ*F/F*_0_)) of individual synapses for correct trials (*n*: *NC* = 518, *WC* = 1555, *SC* = 518 synapses). ***G***, Mean GO Mode, Response and Outcome Direction activity for synapses grouped by coupling. Gray shading indicates the epoch used to calculate activity-direction weights. GO mode (–0.25–0.25 s) projected onto all trial conditions and averaged. Response direction (0–0.6 s) projected onto CR and CL trials and averaged; CL trials multiplied by − 1 to directionally align selectivity. Outcome Direction (0.6–1.2 s) projected onto CR and CL trials and averaged (GO Mode and Outcome Direction *n*: *NC* = 434, *WC* = 1267, *SC* = 479 synapses; Response Direction *n*: *NC* = 518, *WC* = 1555, *SC* = 518 synapses). ***H***, Mean activity in gray windows shown in (G), except GO direction is 0-0.25 s from GO to isolate post-GO activity. For (G-H), error shading indicates hierarchical bootstrap SEM and statistics were calculated with two-tail hierarchical difference test (Table S4).

To quantitatively compare the encoding of task-variables across the coupling groups, we calculated dimensions within the activity space of all synapses (Fig. S5B; 52, 53) that best distinguished the onset of the GO cue (GO Mode), the lick direction (Response Direction), or the task-outcome (Outcome Direction). Projections of all synaptic activity along these dimensions (Fig. S5C) were qualitatively similar to that of local neural activity in ALM cortex (47, 53). Projecting the activity of different coupling groups along these dimensions revealed divergences in task coding (Fig. 4G,H; Fig. S5D; Table S4). The encoding of the GO Mode was greater for *SC* synapses than *NC* synapses (Fig. 4G,H; *SC* = 0.009 ± 0.001 a.u.; *NC* = 0.003 ± 0.001 a.u.; *p* = 0.002), whereas encoding of the Outcome Direction was greater for NC synapses than SC synapses (Fig. 4G,H; *NC* = 0.0026 ± 0.0007; *SC* = 0.0003 ± 0.0001 a.u.; *p* = 6× 10^−4^). The task-encoding of *WC* synapses tended to be intermediate between the two other groups, and Response Direction encoding was similar across groups (Fig. 4G,H).

Next, we investigated whether synaptic activity along these task-related dimensions was associated with the onset of tuft spikes in a manner dependent on coupling group. Despite biases in task encoding, the activity of every coupling group had substantial projections along multiple dimensions (Fig. 4G; Fig. S5D), and thus, a group’s trial-to-trial activity associated with tuft spikes could still be biased to different task-related dimensions than its trial-averaged activity. However, the spike-associated projections of Θ(*dF/dt*)_*TSA*_ along the GO Mode for both *SC*_*SD*_ synapses and *WC*_*SD*_ synapses were greater than for *NC*_*SD*_ synapses (Fig. 5A,B; Table S5; *SC*_*SD*_ = 0.005 ±0.002 a.u.; *WC*_*SD*_ = 0.0021± 0.0008 a.u.; *NC*_*SD*_ = 0.0002 ±0.0003 a.u.; *SC*_*SD*_ vs. *NC*_*SD*_, *p* = 0.003; *WC*_*SD*_ vs. *NC*_*SD*_, *p* = 0.007). In contrast, spike-associated projections along the Outcome Direction were greater for *NC*_*SD*_ synapses than the other two groups (*NC*_*SD*_ = 0.0008 ±0.0005 a.u.; *SC*_*SD*_ = −0.00024± 0.00014 a.u.; *WC*_*SD*_ = −0.0003 ±0.0002 a.u.; *NC*_*SD*_ vs. *SC*_*SD*_, *p* = 0.02; *NC*_*SD*_ vs. *WC*_*SD*_, *p* = 0.02). Projections were dendrite-specific for all coupling groups along the GO Mode (Fig. 5B; Table S5; *SC*_*OD*_ = 0.0007 ±0.0005 a.u.; *WC*_*OD*_ = 0.0004 ±0.0002 a.u.; *NC*_*OD*_ =− 0.0004 ±0.0001 a.u.; *SC*_*SD*_ vs. *SC*_*OD*_, *p* = 0.02; *WC*_*SD*_ vs. *WC*_*OD*_, *p* = 0.01; *NC*_*SD*_ vs. *NC*_*OD*_, *p* = 0.02). Projections along the Outcome direction were not significantly dendrite-specific for any coupling group (Fig. 5D; Table S5), likely reflecting greater noise in the estimation of the outcome mode as a result of low numbers of error trials per session.

**Fig. 5.**
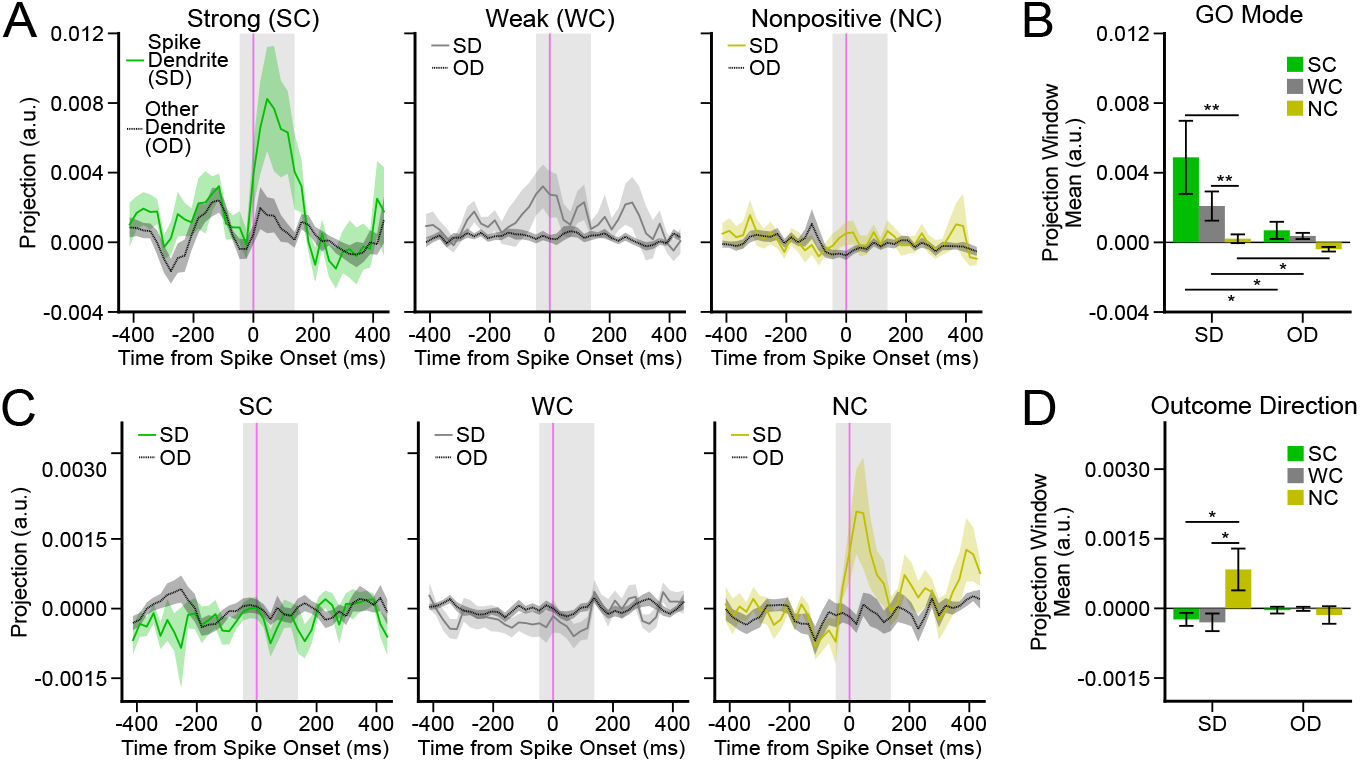
Spike-associated synaptic activity along task-selective dimensions. ***A***, Mean GO Mode projected synaptic activity aligned to tuft spikes by coupling group using Θ(*dF/dt*)_*TSA*_ for SD (solid lines; *n*: *SC*_*SD*_ = 90, *WC*_*SD*_ = 287, *NC*_*SD*_ = 98 synapses) and OD (dashed lines; *n*: *SC*_*OD*_ = 355, *WC*_*OD*_ = 1040, *NC*_*OD*_ = 353 synapses). ***B***, Mean activity in gray windows (–50 to 150 ms from onset) shown in (A). ***C-D***, Same as (A-B), except projected along the Outcome Direction. *n*: *SC*_*SD*_ = 83, *WC*_*SD*_ = 269, *NC*_*SD*_ = 96, *SC*_*OD*_ = 328, *WC*_*OD*_ = 1000, *NC*_*OD*_ = 342. Error shading indicates session-level hierarchical bootstrap SEM and statistics used a one-tailed hierarchical difference test (Table S5).

Taken together, these results suggested that two large-scale networks defined by population coupling encoded different aspects of the task: *SC* inputs preferentially encoded the transition between the preparation epoch and action epoch of the behavioral task, whereas *NC* synapses preferentially encoded task-outcome. These functional differences extended to the spike-associated activity of the synapses coupled to these two networks, and this activity remained dendrite-specific even after partitioning by both task-selectivity and population coupling.

## Discussion

Our results uncover new relationships between spikes in the tuft dendrites and the activity of networks partitioned by coupling and task-coding, providing critical insights into potential roles for dendritic spikes in network-level operations, such as flexible behavior and learning.

Simultaneous high-speed glutamate and calcium imaging allowed us to map the input dynamics accompanying tuft spikes in L5 neurons. Recent studies of synaptic plasticity in layer 2/3 neurons have characterized input timing with respect to bAPs (61–63), but did not investigate trial-to-trial correlations. Thus, our recordings are among the first to precisely measure neuron-specific, trial-to-trial glutamatergic input statistics associated with the generation of postsynaptic events during behavior. We found that tuft spikes were associated with a dendrite-specific increase in input activity within≈ 200 ms of the spike, in addition to the precise onset of additional activity within≈ 70 ms of the spike. Analysis of putative multi-branch Ca^2+^ transients (Fig. S1J,K) and the low frequency of tuft spikes (≈0.01 Hz; Fig. S1C), suggests that most tuft spikes we detected were likely multi-branch regenerative events, and not isolated local dendritic spikes or every bAP. These multi-branch events could be triggered by many mechanisms, including multiple concurrent NMDA spikes, Ca^2+^ spikes originating from the apical nexus, amplified bAPs, or combinations thereof (64).

It is possible that the dendrite-specific increased input associated with tuft spikes reflects shared selectivity between the inputs and the postsynaptic neuron for unmeasured behaviors that varied from trial-to-trial. However, inputs encoding the unmeasured behavior would have to be precisely targeted to individual dendrites to a degree that is difficult to reconcile with existing evidence. Although there is some clustering of inputs by functional selectivity at scales of ≈10 µm (Fig. S2A,B; 19, 60), the tuft dendrites of individual neurons receive inputs with diverse selectivity for behavioral variables across large spatial scales (19).

Alternatively, the dendrite-specific increase in input that accompanies tuft spikes could reflect trial-to-trial variability in the activation of subnetworks that are not exclusively defined by their behavioral selectivity. Both feedforward (65–67) and recurrent (68–73) subnetworks have been proposed as critical circuit mechanisms of signal amplification, persistence, and active filtering. The tuft dendrites of individual L5 ET neurons may receive targeted input from amplifying feedforward subnetworks (67) and cortical-thalamic-cortical loops (74), possibly receiving the output of a chain of loops that traverses the cortical hierarchy (75, 76). Furthermore, recent work has demonstrated that the network influence of individual cortical neurons, rather than their behavioral selectivity, best predicts their influence on behavior (67, 77– 80). Thus, the dendritic spikes and concurrent dendrite-specific increased input we observe may reflect trial-to-trial variable activation of different subnetworks – perhaps due to metastable recurrent dynamics (81–83) – within intermediate dimensions that eventually drive the same action (80). Our results would also be consistent with models in which nonlinear dendritic integration and recurrent activity continuously interact to bind subnetworks (26) or perform learning-related calculations that traverse the cortical hierarchy (28).

Interestingly, we also observed that average tuft-spike-associated input peaked ≈100 ms after the onset of the tuft spike and then persisted over 280 ms (Fig. 2), well beyond the reported kinetics of iGluSnFR3 (rise *<*2 ms; decay = 29 ms; 55). Feedforward inhibition to the distal dendrites of the tuft (84–86) could prevent input late within an input episode from triggering tuft spikes – similar to the temporal gating of action potentials by feedforward inhibition (87–89). Alternatively, this timing could reflect tuft-spike-triggered bursts (1) directly driving looped input (28, 74) or a “prepared state” that facilitates further activity in the subnetwork (26). Furthermore, recent work has found that learning modulates the excitability of the tuft dendrites of specific “engram” L5 ensembles (90).

We uncovered additional evidence suggesting relationships between tuft spikes and network topology by analyzing the population coupling of inputs to the tuft dendrites of L5B neurons. The input population contained a strongly coupled subpopulation, similar to the rich club topology observed in local cortical activity (33–37). Tuft spikes were more precisely aligned to the activity of strongly coupled inputs than weakly coupled inputs, suggesting that the more synchronous activity of strongly coupled inputs may be particularly effective at generating tuft spikes. Surprisingly, simultaneously recorded strongly coupled inputs that did not synapse on the spiking tuft exhibited minimally increased activity (Fig. 3E,F; Fig. 5A,B). Strongly coupled neurons may encompass multiple rich-clubs that are interconnected (36, 91) and these “hub” neurons may provide synchronized – but subnetwork-specific – input to tuft dendrites that is especially effective at triggering tuft spikes. Strongly coupled neurons also exhibit greater plasticity during learning (92–94), changes which our data indicate may be efficiently communicated to the tuft dendrites.

Strongly coupled inputs were particularly active along a “GO” mode (Fig. 4G,H; Fig. 5A,B; Fig. S5D), a dimension that reflects the rapid transition between motor-preparatory and action-ready population activity patterns and that is not selective for trial type (53). Notably, rich-club neurons in layer 2/3 of ALM are also weakly selective for trial-type (37). Our results indicate one possible mechanism by which rich-club activity could trigger rapid population transitions: by facilitating the generation of tuft spikes, which then trigger powerful burst firing (1) or “prepared state” (26) dynamics. Interestingly, we recently found that activity in the tuft of L5 ET neurons in ALM is more precisely aligned to the timing of the GO cue than somatic activity (41), consistent with a potential role for tuft activity in facilitating state transitions.

The anatomical identity of the strongly coupled inputs to the tuft remains unknown. However, converging evidence suggests that thalamocortical inputs play a key role in coordinating a preparation-to-action state transition within ALM (53, 95, 96) as well as state transitions in other cortical regions (97–99). Considering that projections from VM thalamus to ALM cortex preferentially target the apical tuft of L5 ET neurons (74), it is possible that thalamic inputs are over-represented within the strongly coupled population of inputs to the tuft.

Surprisingly, we also found slightly enhanced pairwise correlation between inputs in the lowest quintile of population coupling (Fig. 3D; Fig. S3E) – a topology not unique to synaptic inputs (Fig. S4B) – that preferentially encoded task-outcome (Fig. 4G,H; Fig. 5C,D; Fig. S5D). It is possible these inputs are coupled with previously identified mesoscale cortical networks that distribute emotion information and exhibit similar dynamics (48, 100). Nonpositive coupling synapses also exhibited activity temporally aligned with tuft spikes (Fig. 3E,F), but unlike strong couplers, this activity was along the outcome direction rather than the GO mode (Fig. 5A-D). The outcome information conveyed by nonpositive coupling synapses may facilitate learning-related computations in the tuft (25) or coordinate post-outcome state transitions, similar to the role we propose for strong couplers.

A previous study also reported high noise correlation among outcome-encoding neurons, but did not assess their population coupling (101). Another study (102) reported that weakly coupled neurons were the most selective for gross movement, an inconsistency with our results that may reflect differences between behavioral paradigms and sampling densities used to estimate coupling values.

In summary, we found that tuft spikes are accompanied by temporally extended input episodes that are highly dendrite-specific, suggesting the activation of sparse subnetworks. Yet, the synapses most associated with dendritic spikes belonged either to large-scale coactive input populations that encoded action initiation or one that encoded task outcome. This suggests that tuft spikes may be particularly driven by “hub” neurons that are both embedded in sparse subnetworks and synchronized to large-scale functional networks. Future experiments could employ targeted optogenetics to investigate causal relationships between input coupling and the generation of tuft spikes.

## Code and data availability

Code and data used in the generation of this manuscript are available at https://github.com/kerlin-lab/Gable_2026.

## Acknowledgements

This work was supported with funding from the Whitehall Foundation (Grant 2020-05-71), NINDS R01NS127902, as well as the resources and staff at the University Imaging Centers (UIC; RRID: SCR_020997) and Minnesota Supercomputing Institute (MSI) at the University of Minnesota. Research reported was additionally supported by the University of Minnesota’s MnDRIVE (Minnesota’s Discovery, Research and Innovation Economy) Initiative.

## Author contributions

Designed research: JG, AK. Performed research: JG, SY, JS, SB, NM. Analyzed data: JG, ZN, AK. Wrote the paper: JG. Edited the paper: JG, ZN, AK.

## Supplementary Figures

**Fig. S1.**
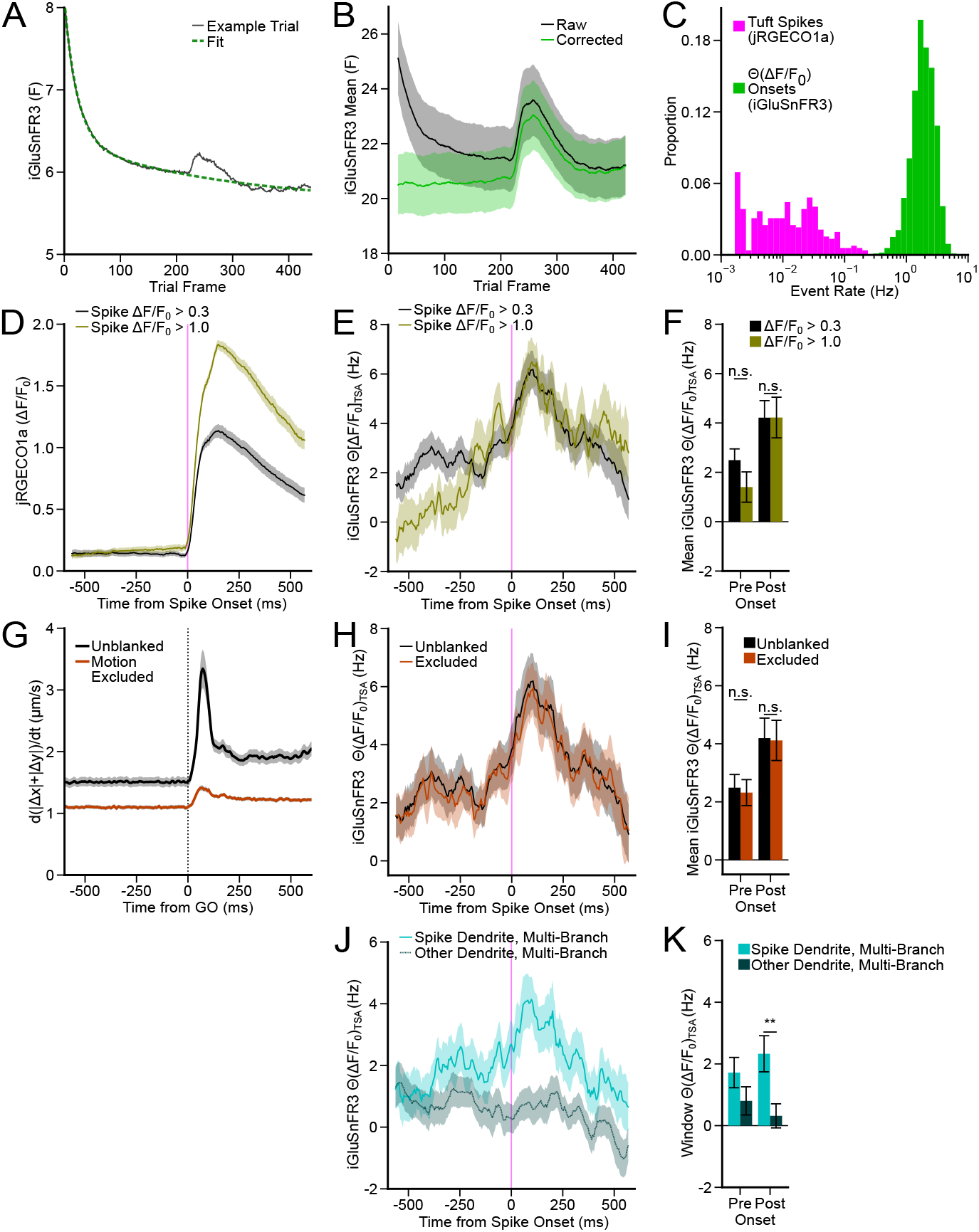
Activity timecourse processing, rates, and robustness. ***A***, Intra-trial photobleaching for example session. Whole-image trace (black) and double exponential decay model fit (dashed green; *τ*_*fast*_ = 23.0 *ms, τ*_*slow*_ = 287 *ms*; *R*^2^ = 0.99). ***B***, Mean of all trial averaged time course of synapses (no pre-processing) with (blue) and without (green) bleach correction for session in (A) (*n* = 64 synapses). ***C***, Event rates of tuft spikes and synaptic Θ(Δ*F/F*_0_) events (*n* = 313 dendrites; *n* = 2072 synapses). ***D***, Mean Ca^2+^ event transient across dendrites using a low (0.3 Δ*F/F*_0_; olive; *n* = 313) and high (1 Δ*F/F*_0_; gray; *n* = 194) minimum event amplitude threshold. ***E***, Same as (D) except mean synaptic activity aligned to calcium transients (Θ(Δ*F/F*_0_)_*T SA*_; low *n* = 826; high *n* = 333). ***F***, Mean pre-onset (–575–0 ms) and post-onset (0–575 ms) activity for data in (E). ***G***, Mean inferred motion (black) and after largest motion frames were removed (dark red; *n* = 52 sessions). ***H***, Mean synaptic Θ(Δ*F/F*_0_) activity aligned to tuft spikes before (black) and after (dark red) removing largest motion frames (*n* = 826 synapses). ***I***, Mean pre-onset (–575–0 ms) and post-onset (0–575 ms) activity for data in (H). ***J***, Average mean-subtracted Θ(Δ*F/F*_0_)_*T SA*_ activity aligned to putative multi-branch tuft spikes. SD synapses (cyan; *n* = 485). OD synapses (gray; *n* = 1332). ***K***, Mean pre-onset (–575–0 ms) and post-onset (0–575 ms) activity for data shown in (J). Error shading and error bars are hierarchical bootstrap SEM. Statistical tests were two-tailed hierarchical difference tests (see Table S6)

**Fig. S2.**
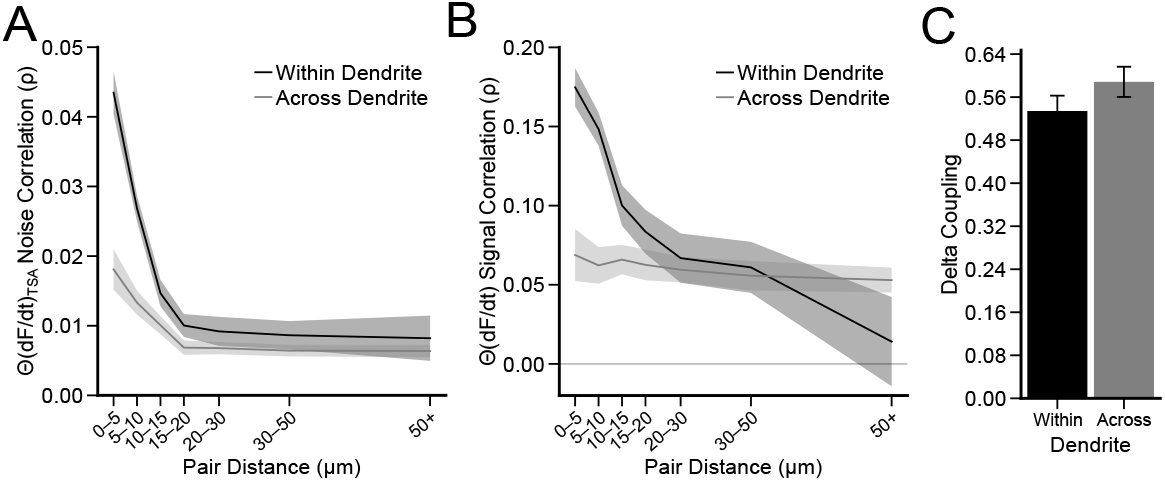
Functional synaptic clustering analysis. ***A***, Mean pairwise synaptic noise correlations (*ρ*) as a function of branch or euclidean distance (Θ(*dF/dt*)_*TSA*_) Pairs located on the same dendrite (black; *n* = 6674 pairs; *R*^2^ = 0.99) or on different dendrites (gray; *n* = 47775 pairs; *R*^2^ = 0.98). ***B***, Same as (A), except for signal correlation (Θ(*dF/dt*) for within dendrite pairs (black; *R*^2^ = 0.96) and across dendrite pairs (gray; *R*^2^ = 0.76). ***C***, Mean pairwise synaptic population coupling similarity for synapses located on the same or different dendrites (*n* = 461 synapses; two-tail hierarchical bootstrap mean difference test *p* = 0.19). Error shading and error bars: hierarchical bootstrap SEM.

**Fig. S3.**
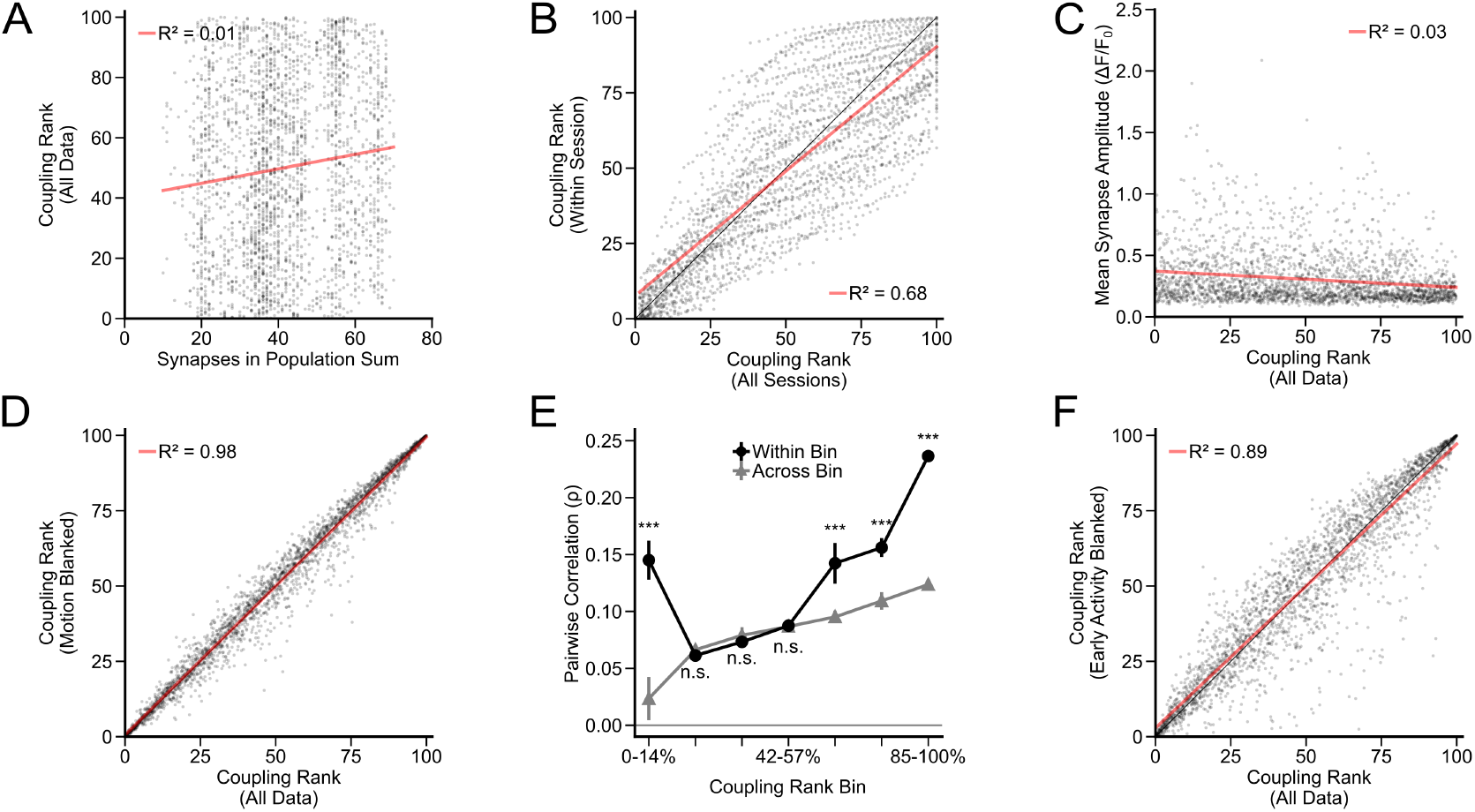
Robustness of population coupling estimates and organization. ***A***, Relationship between coupling rank and population size used to calculate rank. (*r* = 0.12; *n* = 2591 synapses). ***B***, Relationship between synaptic coupling ranks when combining all sessions together versus ranks calculated using single sessions. Red line indicates linear fit (*r* = 0.83; *n* = 2591 synapses) ***C***, Relationship between coupling rank and average individual Δ*F/F*_0_ transient size (*r* = −0.18; *n* = 2591 synapses). ***D***, Relationship between synaptic coupling ranks before and after removing the top 15% of inferred high-motion frames (see Methods for further details). Red line indicates linear fit (*r* = 0.99; *n* = 2591 synapses). ***E***, Mean correlation of synapse pairs by population coupling similarity during spontaneous behavior using Δ*F/F*_0_ traces including pairs in the same coupling range (black circles; *n* = 1965 pairs) and different coupling ranges (gray triangles; *n* = 22424 pairs). Error bars: hierarchical bootstrap SEM. Statistical significance: one-tail hierarchical bootstrap mean difference test (see Table S7). ***F***, Relationship between coupling rank before and after removal of the early response period (0–0.5 s from GO cue) from analysis. Red line indicates linear fit (*r* = 0.94; *n* = 2591 synapses).

**Fig. S4.**
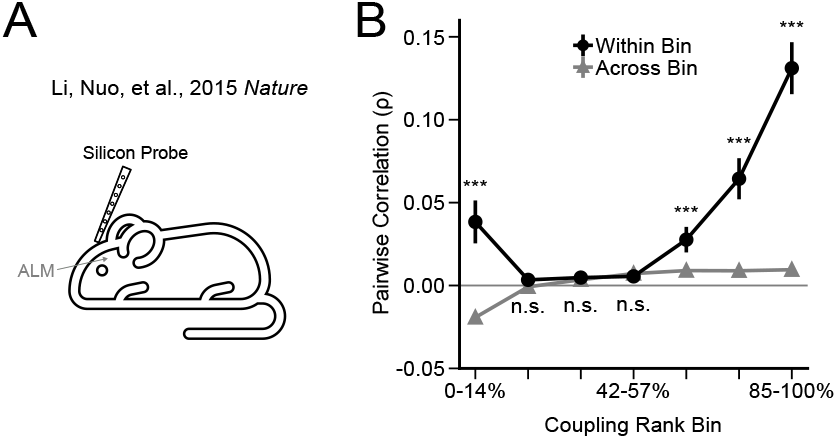
Similar organization of population coupling in silicon probe recordings. ***A***, Secondary data analysis measuring spiking activity at single neuron resolution in ALM using silicon probes in a similar behavior task (49). ***B***, Spiking activity noise correlation (*ρ*) as a function of coupling similarity. Black circles represent pairs in the same coupling range (*n* = 753 pairs). Gray triangles represent pairs in different coupling ranges with one neuron belonging in a given range (*n* = 6586 pairs). Error bars: hierarchical bootstrap SEM. Statistical significance: one-tail hierarchical bootstrap mean difference test (see Table S8).

**Fig. S5.**
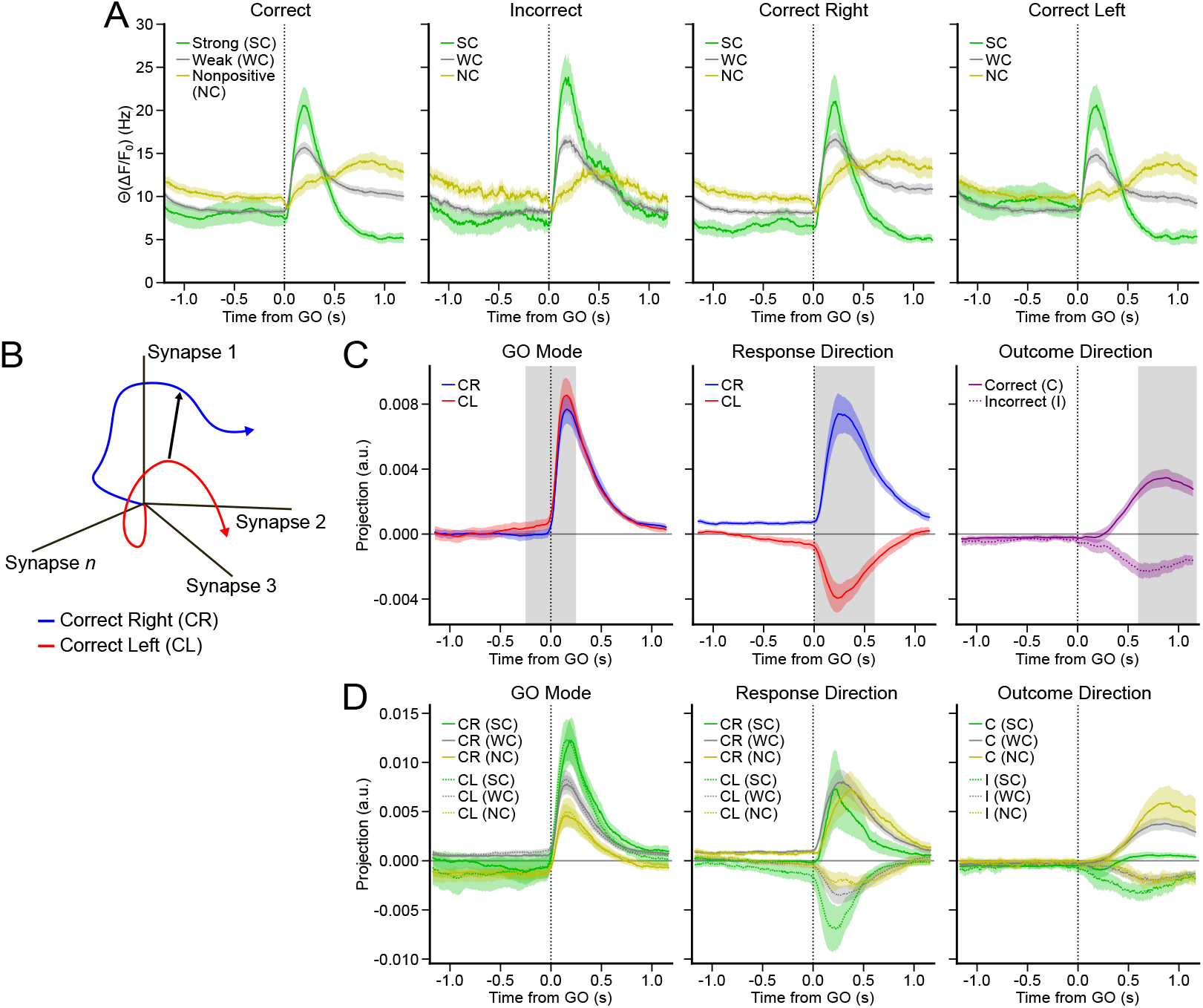
Additional characterization of task coding and population coupling groups. ***A***, Trial averaged mean Θ(Δ*F/F*_0_) aligned to GO cue (dotted black line) for correct (C), incorrect (I), correct right (CR), and correct left (CL) trial conditions for strong (SC), weak (WC) and nonpositive (NC) coupling groups (C/CR/CL *n*: *SC* = 518, *WC* = 1555, *NC* = 518 synapses; I *n*: *SC* = 479, *WC* = 1267, *NC* = 434 synapses). ***B***, Visual depiction of activity mode calculation. ***C***, Population mean for GO mode, Response and Outcome Direction activity for synapses. Θ(Δ*F/F*_0_) traces used for calculation. Gray shading indicates the epoch used to calculate activity-mode weights. GO Mode (–0.25–0.25 s) projected onto CR and CL trials. Response Direction (0–0.6 s) projected onto CR and CL trials. Outcome Direction (0.6–1.2 s) projected onto correct and error trials (GO Mode and Response Direction *n* = 2591 synapses; Outcome Direction *n* = 2180 synapses). ***D***, Same as (C), except split up by population coupling group (GO Mode and Response Direction *n*: *SC* = 518, *WC* = 1555, *NC* = 518; Outcome Direction *n*: *SC* = 479, *WC* = 1267, *NC* = 434 synapses). Error shading: hierarchical bootstrap SEM.

**Table S1.**
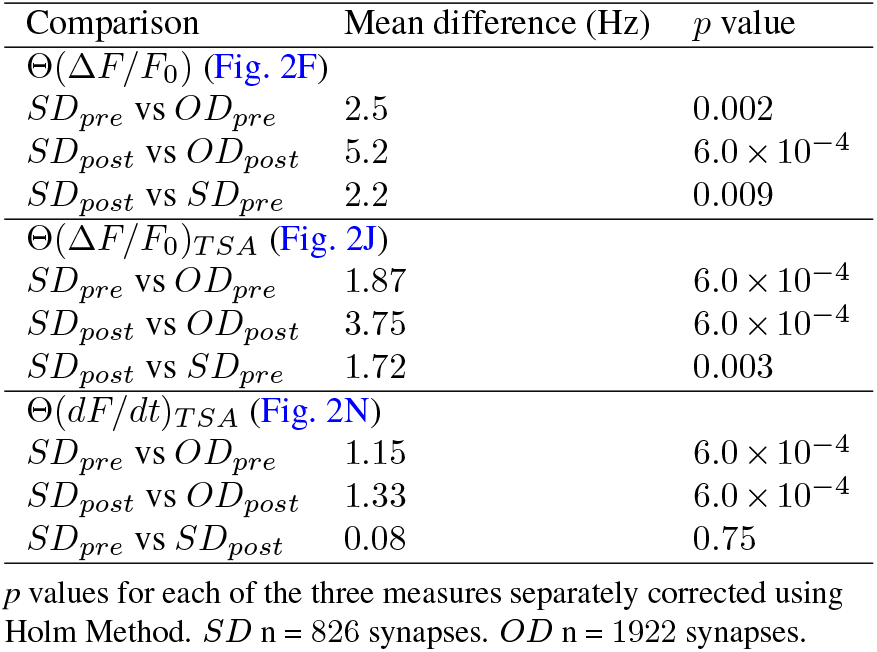
Two-tailed hierarchical bootstrap mean difference tests for population synaptic activity aligned to tuft spikes (Fig. 2).

| Comparison | Mean difference (Hz) | *p* value |
| --- | --- | --- |
| $\Theta(\Delta F/F_0)$ (Fig. 2F) | | |
| $SD_{pre}$ vs $OD_{pre}$ | 2.5 | 0.002 |
| $SD_{post}$ vs $OD_{post}$ | 5.2 | $6.0 \times 10^{-4}$ |
| $SD_{post}$ vs $SD_{pre}$ | 2.2 | 0.009 |
| $\Theta(\Delta F/F_0)_{TSA}$ (Fig. 2J) | | |
| $SD_{pre}$ vs $OD_{pre}$ | 1.87 | $6.0 \times 10^{-4}$ |
| $SD_{post}$ vs $OD_{post}$ | 3.75 | $6.0 \times 10^{-4}$ |
| $SD_{post}$ vs $SD_{pre}$ | 1.72 | 0.003 |
| $\Theta(dF/dt)_{TSA}$ (Fig. 2N) | | |
| $SD_{pre}$ vs $OD_{pre}$ | 1.15 | $6.0 \times 10^{-4}$ |
| $SD_{post}$ vs $OD_{post}$ | 1.33 | $6.0 \times 10^{-4}$ |
| $SD_{pre}$ vs $SD_{post}$ | 0.08 | 0.75 |

*p* values for each of the three measures separately corrected using Holm Method. *SD* *n* = 826 synapses. *OD* *n* = 1922 synapses.

**Table S2.** One-tailed hierarchical bootstrap mean difference tests for same and different coupling range pairwise noise correlations (Fig. 3D).

| Bin | Mean difference ($\rho$) | *p* value |
| --- | --- | --- |
| 0.00%–14.29% | 0.02 | $7.0 \times 10^{-3}$ |
| 14.29%–28.57% | 0.002 | 0.11 |
| 28.57%–42.86% | 0.0007 | 0.24 |
| 42.86%–57.14% | 0.002 | 0.11 |
| 57.14%–71.43% | 0.01 | $7.0 \times 10^{-3}$ |
| 71.43%–85.71% | 0.02 | $7.0 \times 10^{-3}$ |
| 85.71%–100.00% | 0.06 | $7.0 \times 10^{-3}$ |

*p* values corrected using Holm method. Same coupling range *n* = 9561 pairs. Different coupling range *n* = 71212 pairs.

**Table S3.** Two-tailed hierarchical bootstrap mean difference tests for synaptic activity aligned to tuft spikes by coupling group (Fig. 3F).

| Comparison | Mean difference (Hz) | *p* value |
| --- | --- | --- |
| $SC_{SD}$ vs $WC_{SD}$ | 1.1 | 0.002 |
| $SC_{SD}$ vs $NC_{SD}$ | 1.0 | 0.02 |
| $NC_{SD}$ vs $WC_{SD}$ | 0.08 | 0.7 |
| $SC_{SD}$ vs $SC_{OD}$ | 1.96 | $6.0 \times 10^{-4}$ |
| $WC_{SD}$ vs $WC_{OD}$ | 0.59 | $6.0 \times 10^{-4}$ |
| $NC_{SD}$ vs $NC_{OD}$ | 0.77 | $6.0 \times 10^{-4}$ |

Between-coupler-group *SD*, and *SD* vs *OD* *p* values separately corrected using Holm method. $NC_{SD}$ *n* = 160 synapses, $NC_{OD}$ *n* = 384, $WC_{SD}$ *n* = 522, $WC_{OD}$ *n* = 1156, $SC_{SD}$ *n* = 144, $SC_{OD}$ *n* = 382.

**Table S4.**
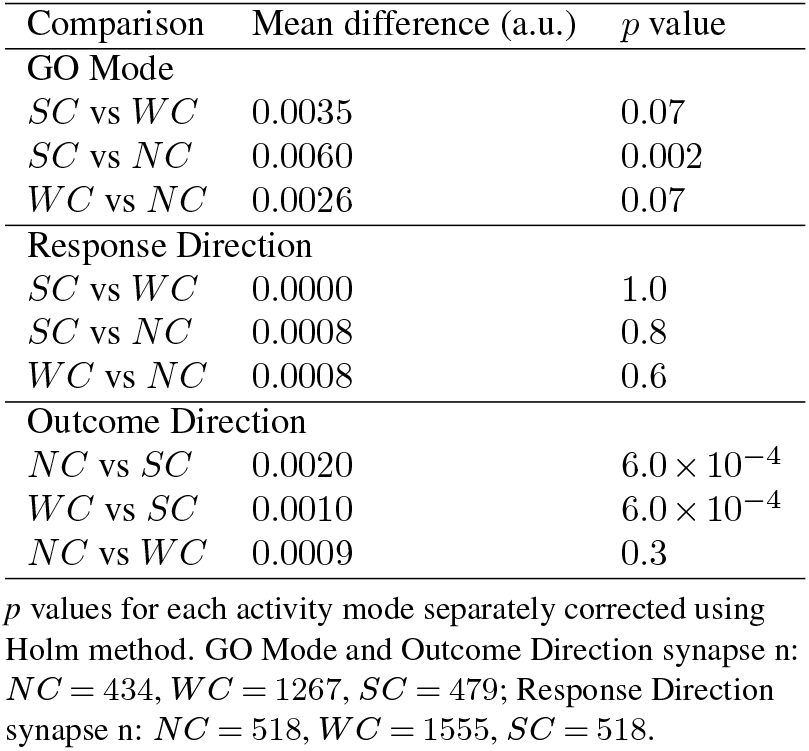
Two-tailed hierarchical difference tests for synaptic activity modes aligned to GO cue. (Fig. 4H).

**Table S5.** One-tailed hierarchical difference tests for GO Mode and Outcome Direction synaptic activity aligned to tuft spikes (Fig. 5).

| Comparison | Mean difference (a.u.) | *p* value |
| --- | --- | --- |
| **GO Mode (Fig. 5B)** |  |  |
| *SC<sub>SD</sub>* vs *WC<sub>SD</sub>* | 0.003 | 0.1 |
| *SC<sub>SD</sub>* vs *NC<sub>SD</sub>* | 0.005 | 0.003 |
| *WC<sub>SD</sub>* vs *NC<sub>SD</sub>* | 0.002 | 0.007 |
| *SC<sub>SD</sub>* vs *SC<sub>OD</sub>* | 0.004 | 0.02 |
| *WC<sub>SD</sub>* vs *WC<sub>OD</sub>* | 0.002 | 0.01 |
| *NC<sub>SD</sub>* vs *NC<sub>OD</sub>* | 0.0006 | 0.02 |
| **Outcome Direction (Fig. 5D)** |  |  |
| *NC<sub>SD</sub>* vs *SC<sub>SD</sub>* | 0.001 | 0.02 |
| *NC<sub>SD</sub>* vs *WC<sub>SD</sub>* | 0.001 | 0.02 |
| *WC<sub>SD</sub>* vs *SC<sub>SD</sub>* | 0 | 0.6 |
| *NC<sub>SD</sub>* vs *NC<sub>OD</sub>* | 0.001 | 0.05 |
| *WC<sub>SD</sub>* vs *WC<sub>OD</sub>* | -0.0003 | 1 |
| *SC<sub>SD</sub>* vs *SC<sub>OD</sub>* | -0.0002 | 1 |

*p* values separately corrected using Holm method for each group of hypotheses (GO Mode SD across coupler groups, GO Mode SD vs OD, Outcome Direction SD across coupler groups, Outcome Direction SD vs OD). GO Mode synapse n: *SC<sub>SD</sub>* = 90, *WC<sub>SD</sub>* = 287, *NC<sub>SD</sub>* = 98, *SC<sub>OD</sub>* = 355, *WC<sub>OD</sub>* = 1040, *NC<sub>OD</sub>* = 353. Outcome Direction synapse n: *SC<sub>SD</sub>* = 83, *WC<sub>SD</sub>* = 269, *NC<sub>SD</sub>* = 96, *SC<sub>OD</sub>* = 328, *WC<sub>OD</sub>* = 1000, *NC<sub>OD</sub>* = 342.

**Table S6.**
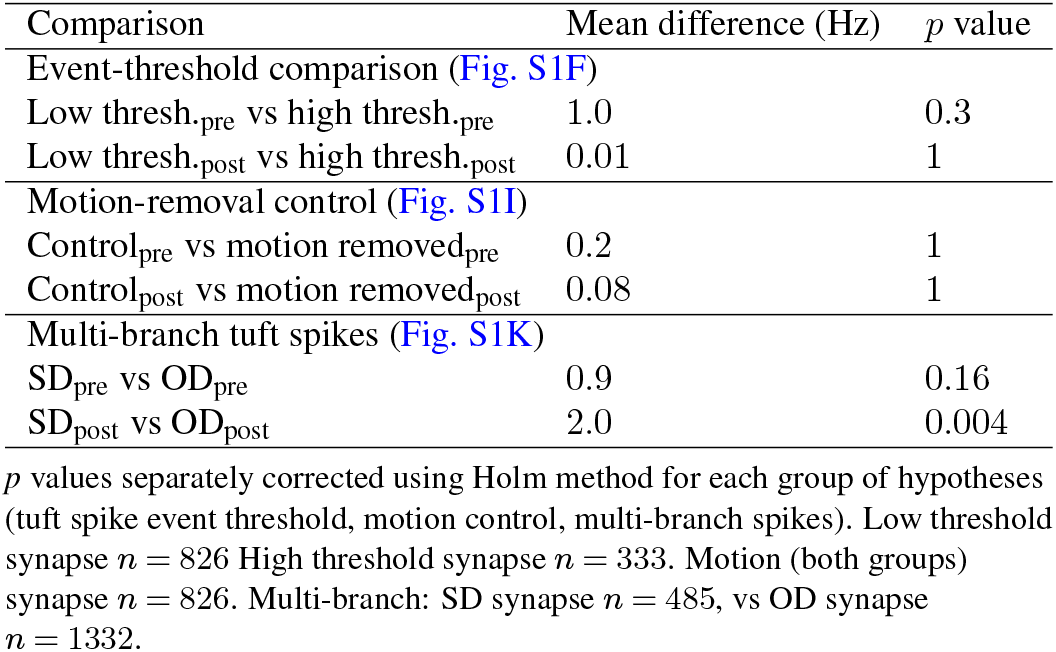
Two-tailed hierarchical bootstrap mean difference tests for tuft spike-associated synaptic activity control analyses (Fig. S1).

| Comparison | Mean difference (Hz) | *p* value |
| --- | --- | --- |
| Event-threshold comparison (Fig. S1F) |  |  |
| Low thresh.<sub>pre</sub> vs high thresh.<sub>pre</sub> | 1.0 | 0.3 |
| Low thresh.<sub>post</sub> vs high thresh.<sub>post</sub> | 0.01 | 1 |
| Motion-removal control (Fig. S1I) |  |  |
| Control<sub>pre</sub> vs motion removed<sub>pre</sub> | 0.2 | 1 |
| Control<sub>post</sub> vs motion removed<sub>post</sub> | 0.08 | 1 |
| Multi-branch tuft spikes (Fig. S1K) |  |  |
| SD<sub>pre</sub> vs OD<sub>pre</sub> | 0.9 | 0.16 |
| SD<sub>post</sub> vs OD<sub>post</sub> | 2.0 | 0.004 |

*p* values separately corrected using Holm method for each group of hypotheses (tuft spike event threshold, motion control, multi-branch spikes). Low threshold synapse *n* = 826 High threshold synapse *n* = 333. Motion (both groups) synapse *n* = 826. Multi-branch: SD synapse *n* = 485, vs OD synapse *n* = 1332.

**Table S7.** One-tailed hierarchical mean difference tests for same and different coupling range pairwise correlations during spontaneous behavior (Fig. S3E).

| Bin | Mean difference ($\rho$) | *p* value |
| --- | --- | --- |
| 0.00%–14.29% | 0.1 | $7.0 \times 10^{-4}$ |
| 14.29%–28.57% | -0.005 | 1 |
| 28.57%–42.86% | -0.006 | 1 |
| 42.86%–57.14% | 0.0007 | 1 |
| 57.14%–71.43% | 0.05 | $7.0 \times 10^{-4}$ |
| 71.43%–85.71% | 0.05 | $7.0 \times 10^{-4}$ |
| 85.71%–100.00% | 0.1 | $7.0 \times 10^{-4}$ |

*p* values corrected using Holm method. Mice *n* = 1. Session *n* = 3. Same coupling range *n* = 1965 pairs. Different coupling range *n* = 22424 pairs.

**Table S8.**
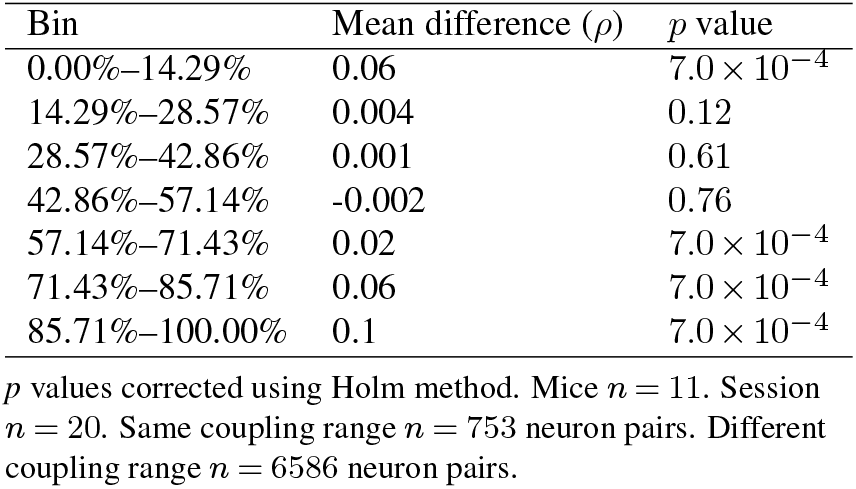
One-tailed hierarchical mean difference tests for same and different coupling range pairwise spiking correlations at somas in ALM (Fig. S4B).

